# Actin reorganization at the centrosomal area and the immune synapse regulates polarized secretory traffic of multivesicular bodies in T lymphocytes

**DOI:** 10.1101/2020.05.01.071712

**Authors:** Ana Bello-Gamboa, Marta Velasco, Solange Moreno, Gonzalo Herranz, Roxana Ilie, Silvia Huetos, Sergio Dávila, Alicia Sánchez, Jorge Bernardino de la Serna, Víctor Calvo, Manuel Izquierdo

## Abstract

T-cell receptor stimulation induces the convergence of multivesicular bodies towards the microtubule-organizing center (MTOC) and the polarization of the MTOC to the immune synapse (IS). These events lead to exosome secretion at the IS. We describe here that upon IS formation centrosomal area F-actin decreased concomitantly with MTOC polarization to the IS. PKCδ-interfered T cell clones showed a sustained level of centrosomal area F-actin associated with defective MTOC polarization. We analysed the contribution of two actin cytoskeleton-regulatory proteins, FMNL1 and paxillin, to the regulation of cortical and centrosomal F-actin networks. FMNL1β phosphorylation and F-actin reorganization at the IS were inhibited in PKCδ-interfered clones. F-actin depletion at the central region of the IS, a requirement for MTOC polarization, was associated with FMNL1β phosphorylation at its C-terminal, autoregulatory region. Interfering all FMNL1 isoforms prevented MTOC polarization; nonetheless, FMNL1β re-expression restored MTOC polarization in a centrosomal area F-actin reorganization-independent manner. Moreover, PKCδ-interfered clones exhibited decreased paxillin phosphorylation at the MTOC, which suggests an alternative actin cytoskeleton regulatory pathway. Our results infer that PKCδ regulates MTOC polarization and secretory traffic leading to exosome secretion in a coordinated manner by means of two distinct pathways, one involving FMNL1β regulation and controlling F-actin reorganization at the IS, and the other, comprising paxillin phosphorylation potentially controlling centrosomal area F-actin reorganization.

## INTRODUCTION

T cell receptor (TCR) stimulation by antigen presented by major histocompatibility complex (MHC) molecules on an antigen-presenting cell (APC) induces the formation of the immune synapse (IS), the convergence of secretory vesicles of T lymphocytes towards the microtubule-organizing center (MTOC) and the polarization of the MTOC to the IS [1, 2]. IS formation is associated with an initial increase in cortical actin at the IS [3]. Subsequently, MTOC polarization is preceded by a decrease in cortical actin density at the central region of the immune synapse (cIS), that contains the secretory domain [4] [5]. The central supramolecular activation cluster (cSMAC) is located in this low-density F-actin region of the cIS (Fact-Low cIS), adjacent to the secretory domain [2, 4, 5]. Next, cortical actin recovery at the IS leads to termination of lytic granule secretion in CTLs [6]. These reversible, cortical actin cytoskeleton reorganization processes occur during the degranulation of both the lytic granules in cytotoxic T lymphocytes (CTL) and some cytokine-containing secretory vesicles in T-helper (Th) lymphocytes [4, 5, 7], despite both the nature and cargo of the secretory vesicles in these cell types are quite different. Thus, the formation of a mature IS produces several crucial biological outcomes, including activation of Th cells (usually CD4^+^ cells) and cell killing by primed CTLs (mostly CD8+ cells). Accordingly, there are two major types of secretory IS established by T lymphocytes that result in very diverse, but also critical, immune effector functions [8] [9] [10]. The IS made by primed CTLs induces rapid polarization (ranging from seconds to few minutes) of the lytic granules towards the IS. The degranulation of the lytic granules induces the secretion perforin and granzymes to the synaptic cleft [11], which are pro-apoptotic molecules, but also exosomes [12]. The secreted perforin and granzymes subsequently induce killing of the target cells [13]. CTLs develop transient synapses, lasting only few minutes, as the target cells are killed [4] [2]. This is probably due to the fact that the optimal CTL task requires a rapid and temporary contact with target cells, in order to distribute as many lethal strikes as possible to numerous target cells [4] [2]. In contrast, Th lymphocytes such as Jurkat cells generate stable, long-standing IS (from 10-30 min up to hours), since this appears to be necessary for both directional and continuous secretion of stimulating cytokines [4] [2]. Moreover, we have described in Th lymphocytes that cortical actin reorganization at the IS plays an important role during the polarized traffic of multivesicular bodies (MVB) containing intraluminal vesicles (ILV) [14], a different type of secretory vesicles involved in exosome secretion at the IS [15–17]. Exosome secretion in Th lymphocytes follows TCR stimulation, IS formation, MVB polarization and degranulation, and these exosomes can induce Fas ligand-dependent autocrine activation-induced cell death (AICD) [18] [16] [15], a key process in regulating T lymphocyte homeostasis [19]. Regarding the intracellular signals controlling this specialized secretory pathway, we have shown that TCR-stimulated PKCδ regulates cortical actin reorganization at the IS ultimately controlling MTOC/MVB polarization leading to exosome secretion at the IS [14]. PKCδ-interfered T cell clones showed inhibition of cortical actin reorganization at the IS concomitantly to defective MTOC/MVB polarization [14]. Overall, this lead us to hypothesize that an altered actin reorganization at the IS may underlie the deficient MVB polarization occurring in PKCδ-interfered T cell clones [14]. However, it cannot be ruled out that PKCδ may govern other actin reorganization networks apart from actin reorganization at the IS, that may also contribute to the diminished MTOC/MVB polarization efficiency observed in PKCδ-interfered clones [14].

TCR-triggered actin cytoskeleton reorganization and polarized traffic of secretory vesicles are regulated by two major pathways: one involves HS1/WASp/Arp2/3 complexes acting in cortical F-actin, and the other formins such as formin-like 1 (FMNL1) and Diaphanous-1 (Dia1) [20] [21]. Several studies suggest that cortical F-actin reorganization at the IS is necessary and sufficient for MTOC/lytic granules polarization [5] [22]. However, other results show that depletion of formins FMNL1 or Dia1 impedes MTOC polarization without affecting Arp2/3-dependent cortical actin reorganization [23], supporting that, at least in the absence of FMNL1 or Dia1, cortical actin reorganization is not sufficient for MTOC polarization. Conversely, in the absence of cortical actin reorganization at the IS occurring in Jurkat cells lacking Arp2/3, the MTOC can polarize normally to the IS [23] [21]. Regarding other molecules controlling MTOC polarization, paxillin (an actin-regulating adaptor protein) [24] is necessary for MTOC polarization to the IS in CTLs [25] and its phosphorylation is required for lytic granule secretion from CTL [26]. Along these lines, recent studies in B lymphocytes forming IS indicated that the MTOC is a F-actin organizing center [27], and F-actin depletion around the MTOC crucially facilitates MTOC polarization towards the IS in B-cell receptor (BCR)-stimulated B lymphocytes [28] [29]. In summary, taking together these observations, it is clear that MTOC and MVB polarization induced by IS formation are regulated by HS1/WASp/Arp2/3- dependent cortical and formin-dependent non-cortical actin networks, and that PKCδ may regulate the reorganization of both actin networks.

To better understand the mechanisms involved in these processes in T cells, we studied the contribution of FMNL1, Dia1 and paxillin. Here we report that PKCδ appears to coordinately regulate MTOC and MVB polarizations via at least two distinct actin cytoskeleton regulatory pathways: FMNL1β-mediated regulation of F-actin reorganization at the IS and paxillin-mediated reorganization of centrosomal area F-actin.

## MATERIALS AND METHODS

### Cells

Raji B and Jurkat T (clone JE6.1) cell lines were obtained from the ATCC. Cell lines were cultured in RPMI 1640 medium containing L-glutamine (Invitrogen) with 10% heat-inactivated FCS (Gibco) and penicillin/streptomycin (Gibco). Jurkat cells were transfected with control or PKCδ shRNA-encoding plasmids, selected with puromycin, and control (C3, C9) and PKCδ-interfered (P5, P6) Jurkat stable clones were isolated by limiting dilution and phenotyped as previously described [14].

### Plasmids and transient transfection

pECFP-C1CD63 (expressing CFP-CD63) was provided by G. Griffiths. CD63 is an enriched marker in MVB and has been widely used to study the secretory traffic of MVB both in living and fixed cells [15, 16]. The plasmid expressing dsRed-centrin 2 (dsRed-cent2) was a gift from J. Gleeson (Addgene plasmid # 29523; http://n2t.net/addgene:29523; RRID:Addgene_29523). pmCherry Paxillin was a gift from K. Yamada (Addgene plasmid # 50526; http://n2t.net/addgene:50526; RRID:Addgene_50526). The control vector (shControl-HA-YFP), FMNL1 interfering (shFMNL1-HA-YFP) and FMNL1-interfering, re-expressing vectors (shFMNL1-HA-YFP-FMNL1α, shFMNL1-HA-YFP-FMNL1β, shFMNL1-HA-YFP-FMNL1γ and shFMNL1-HA-YFP-FMNL1ΔFH2 mutant) were previously described [30] and generously provided by D. Billadeau. Jurkat clones were transiently transfected with 20-30 μg of the plasmids as described [16].

### Antibodies and reagents

Rabbit monoclonal anti-human PKCδ EP1486Y for WB, does not recognize mouse PKCδ (Abcam). Rabbit monoclonal anti-PKCδ EPR17075 for WB recognizes both human and mouse PKCδ (Abcam). Mouse monoclonal anti-human CD3 UCHT1 for cell stimulation and immunofluorescence (BD Biosciences and Santa Cruz Biotechnology). Mouse monoclonal anti-FMNL1 clone C-5 for WB, and mouse monoclonal anti-FMNL1 clone A-4 for immunoprecipitation (Santa Cruz Biotechnology). Mouse monoclonal anti-Dia1 clone E-4 for WB (Santa Cruz Biotechnology) and rabbit polyclonal anti-Dia1 (Diap1) clone E1E4K for immunoprecipitation (Cell Signalling Technology). Mouse monoclonal anti-γ−tubulin for immunofluorescence (SIGMA). Mouse monoclonal anti-paxillin clone 349 for WB and anti-paxillin clone 349 coupled to TRITC for immunofluorescence (BD Biosciences). Rabbit polyclonal anti-phospho-Thr538 paxillin for WB and immunofluorescence (ECM Biosciences). Rabbit polyclonal Phospho-(Ser) PKC substrate antibody for WB and immunofluorescence (Cell signalling Technology). Fluorochrome-coupled secondary antibodies (goat-anti-mouse IgG AF488 A-11029, goat-anti-rabbit IgG AF488 A-11034, goat-anti-mouse IgG AF546 A-11030, goat-anti-mouse IgG AF647 A-21236) for immunofluorescence were from ThermoFisher. Horseradish peroxidase (HRP)-coupled secondary antibodies (goat anti-mouse IgG-HRP, sc-2005 and goat anti-rabbit IgG-HRP, sc-2004) were from Santa Cruz Biotechnology. CellTracker™ Blue (CMAC) and phalloidin were from ThermoFisher. Staphylococcal enterotoxin E (SEE) was from Toxin Technology, Inc. SirActin and verapamil were from Cytoskeleton Inc.

### Immunoprecipitation

Immunoprecipitation from cell lysates was performed by using Protein A/G Magnetic Beads (Pierce, ThermoScientific) following the instructions provided by the company. Briefly, 0.5 ml lysates corresponding to 20-30×10^6^ Jurkat clones, stimulated or not as described, were incubated with the primary antibody (5 μg) for 2 h at 4° C. Subsequently, 15 μl of magnetic beads suspension were added and incubated for 3 h at 4° C. Beads were washed 5x with lysis buffer and the antigens were eluted with 2 M glycine pH=2 and neutralized. Eluates were run on SDS-PAGE gels, 6,5 % Acrylamide and proteins transferred to PDVF membranes.

### Western blot analysis

Cells were lysed in Triton^TM^ x100-containing lysis buffer supplemented with both protease and phosphatase inhibitors. Approximately 50 μg of cellular proteins were recovered in the 10,000xg pellet from 10^6^ cells. Cell lysates and neutralized, acid-eluted immunoprecipitates were separated by SDS-PAGE under reducing conditions and transferred to Hybond^TM^ ECL^TM^ membranes (GE Healthcare). Membranes were incubated sequentially with the different primary antibodies and developed with the appropriate HRP-conjugated secondary antibody using enhanced chemiluminescence (ECL). When required, the blots were stripped following standard protocols prior to reprobing them with primary and HRP-conjugated secondary antibodies. Autoradiography films were scanned and the bands were quantified using Quantity One 4.4.0 (Bio-Rad) and ImageJ (Rasband, W.S., ImageJ, National Institutes of Health, Bethesda, Maryland, USA, http://rsb.info.nih.gov/ij/, 1997-2004) softwares.

### Time-lapse microscopy, immunofluorescence and image analysis

Jurkat clones transfected with the different expression plasmids were attached to ibidi microwell culture dishes using fibronectin (0.1 mg/ml) at 24-48 h post-transfection, and stimulated in culture medium at 37 °C. In some experiments requiring IS formation, Raji cells attached to ibidi microwell culture dishes using fibronectin (ibiTreat, for paraformaldehyde fixing) or poly-L-lysine (glass bottom, for acetone-fixation) were labelled with CMAC and pulsed with 1 μg/ml SEE, mixed with transfected Jurkat clones and immune synapses analyzed as described [31] [15] [32]. Acetone fixation required for γ-tubulin staining of MTOC was compatible with phalloidin labeling [33]. In other experiments, transfected Jurkat clones were stimulated with plastic-bound anti-TCR UCHT1 (10 μg/ml) or in suspension with phorbol myristate acetate (PMA, 100 ng/ml). Immunofluorescence of fixed synapses was performed as previously described [34], and additional fixations were performed between each fluorochrome-coupled secondary antibody staining and subsequent fluorochrome-coupled primary antibody staining, to exclude any potential cross-reaction of secondary antibodies (i.e. Fig. 7A).

For in vivo actin reorganization experiments, dsRed-Cent2-transfected Jurkat clones were preincubated overnight with 100 nM SirActin and 2 μM verapamil, and subsequently challenged with SEE-pulsed Raji cells as described above. Wide-field, time-lapse micorscopy was performed using an OKO-lab stage incubator (OKO) on a Nikon Eclipse TiE microscope equipped with a DS-Qi1MC digital camera and a PlanApo VC 60x/1.4NA OIL objective (Nikon). Time-lapse acquisition and analysis were performed by using NIS-AR software (Nikon). Subsequently, epi-fluorescence images were improved by Huygens Deconvolution Software from Scientific Volume Image (SVI) using the “widefield” optical option as previously described [35] [32]. For quantification, digital images were analyzed using NIS-AR (Nikon) or ImageJ softwares (Rasband, W.S., ImageJ, National Institutes of Health, Bethesda, Maryland, USA, http://rsb.info.nih.gov/ij/, 1997-2004). The quantification and analysis of F-actin mean fluorescence intensity (MFI) in a centrosome-centered area (centrosomal area F-actin MFI) in time-lapse experiments, was performed within a 2 µm diameter, floating region of interest (ROI) (i.e., ROI changing XY position over time), centered at the center of mass of the MTOC (MTOC^c^), by using NIS-AR software (Fig. 2D). These measurements were performed in deconvoluted time-lapse series because of the enhanced signal-to-noise ratio of the images, although raw time-lapse series yielded comparable results. In parallel, for each time-lapse time point, the measurement of the distance from the MTOC^c^ towards the IS was performed by using NIS-AR software and represented versus the corresponding centrosomal area F-actin MFI value (Fig. 2D, upper panels). Confocal microscopy imaging of synapses made by living cells was performed by using a SP8 Leica confocal microscope equipped with an HC PL APO CS2 63x/1.2 NA water objective (zoom for C3, 5.34; zoom for P5, 4.47; scan velocity, 1000 Hz bidirectional; pixel size, 0.068 μm; pinhole, 111.5 μm; z-step size, 0.6 μm; z-stack, 9 μm). The synaptic conjugates for these experiments were prepared by mixing SEE-pulsed Raji cells with transfected Jurkat clones in suspension, as previously described [5, 6]. The quantification of relative centrosomal area F-actin MFI in these experiments was calculated as the F-actin MFI corresponding to a 2 µm diameter, floating ROI, centered at the MTOC center of mass (MTOC^c^), relative to the F-actin MFI of this centrosomal area ROI at time=0, using the average intensity projection (AIP) from the three focal planes (2 µm thickness) containing the maximal signal of MTOC.

Confocal microscopy imaging in fixed synapses was performed by using a SP8 Leica confocal microscope, with sequential acquisition, bidirectional scanning and the following laser lines: UV (405 nm, intensity: 33.4%), supercontinuum visible (633 nm, intensity: 15.2%), supercontinuum visible (550 nm, intensity: 20.8%), supercontinuum visible (488 nm, intensity: 31.2%). Deconvolution of confocal images was performed by using Huygens Deconvolution Software from Scientific Volume Image (SVI) with the “confocal” optical option. Colocalization analyses were accomplished by using Jacop plugin from ImageJ.

The velocity of movement of MVB was measured by automatically analyzing the trajectories of CFP-CD63^+^ vesicles in videos (i.e. Suppl. Video 2) with the use of NIS-AR software (tracking module) and the ImageJ MJTrack plugin. The trajectory of MTOC was analysed by using ImageJ MJTrack plugin (i.e. Suppl. Video 1). In polarization experiments, to establish the relative ability of the MTOC and MVB to polarize towards the IS, MTOC and MVB polarization indexes (Pol. Indexes) were calculated as described in Fig. 1A, using MIP of acetone-fixed synapses. In the MIP, the position of the cell center of mass (Cell^C^), MTOC and MVB center of mass (MTOC^C^ and MVB^C^, respectively) were used to project MTOC^C^ (or MVB^C^) on the vector defined by the Cell^C^–synapse axis. Then the MTOC (or MVB) polarization index was calculated by dividing the distance between the MTOC^C^ (or MVB^C^) projection and the Cell^C^ (“A” distance) by the distance between the Cell^C^ and the synapse (“B” distance) (Fig. 1A). Cell^C^ position was taken as the origin to measure distances, thus those “A” values in the opposite direction to the synapse were taken as negative. Thus, Pol. Indexes (Pol. Index = A/B) ranked from +1 (fully polarized) to -1 (fully anti-polarized). Therefore Pol. Index values were normalized by cell size and shape (Fig. 1A). For the experiments analyzing F-actin reorganization at MTOC area in fixed cells, the MTOC area-associated F-actin (centrosomal area F-actin) was quantified by labelling MTOC with γ-tubulin or expressing dsRed-Cent2, since both labelling approaches rendered equivalent results, as shown in Fig. 2 and Suppl. Fig. S3. Briefly, after manual selection of the z optical section containing the maximal signal corresponding to the centrosome (γ-tubulin or dsRed-Cent2 signal), a substack (2 μm thickness) centered at the MTOC^c^ was selected, and the F-actin average intensity z-projection (AIP) of the substack was generated (centrosomal area AIP) and thresholded (Default), by using ImageJ. Subsequently, a circular ROI “C” (2 μm diameter) centered at the MTOC^c^ was selected on the thresholded, centrosomal area AIP, and the F-actin MFI in this ROI was calculated, and this value corresponded to centrosomal area F-actin MFI (MFI in ROI “C”, Fig. 2A and Suppl. Fig. S3). In parallel, we calculated the cellular F-actin MFI of the thresholded (Default) average intensity z-projection (AIP) corresponding to all focal planes (total AIP), using a ROI including the whole cell (MFI in ROI “D”, Fig. 2A and Suppl. Fig. S3). We then calculated the centrosomal area F-actin MFI ratio (centrosomal area F-actin MFI/ cell F-actin MFI= MFI C/MFI D) to normalize by cell size and by phalloidin labelling among different samples. This value represented the relative density of centrosomal area F-actin (Fig. 2, Suppl. Fig. S3). For paxillin phosphorylation image analyses, image quantification of phospho-T538 MFI signal in fixed synapses was performed by using ROI defined with the “autodetect” algorithm from NIS-AR, containing the paxillin fluorescence signal. These phospho-T538 MFI values were internally normalized by paxillin MFI values in the same ROI by using ImageJ. For image analyses of FMNL1 isoform phosphorylation, image quantification of Phospho-(Ser) PKC substrate signal was performed by using cell ROIs defined with the “Autodetect” algorithm from NIS-AR and an appropriate threshold. Image analysis data correspond to at least three different experiments, analyzing a minimum of 30 synapses from 15 different, randomly selected, microscopy fields per experiment. ANOVA analysis was performed for statistical significance of the results using Excel and IBM’s SPSS Statistics software.

**Fig. 1.**
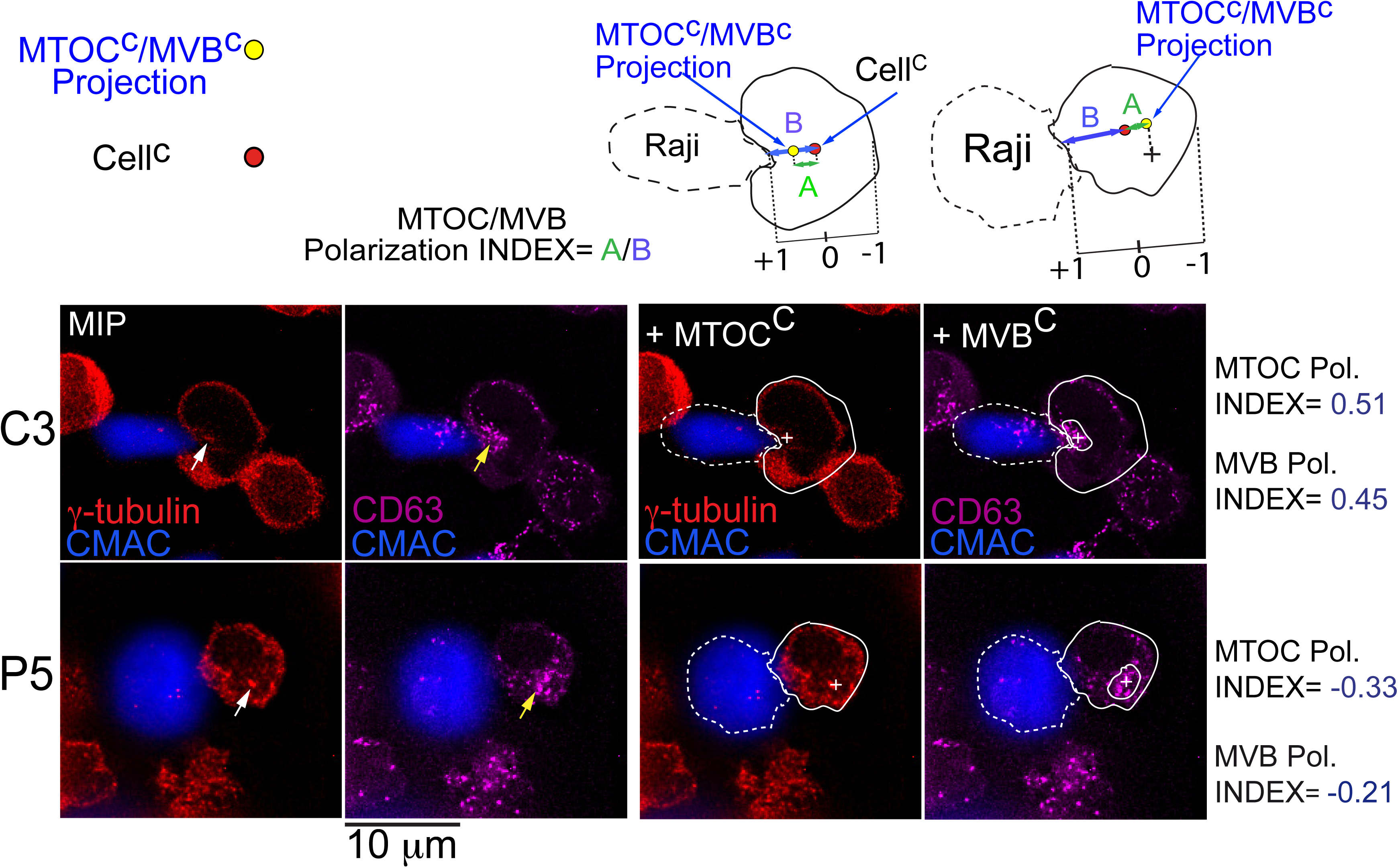
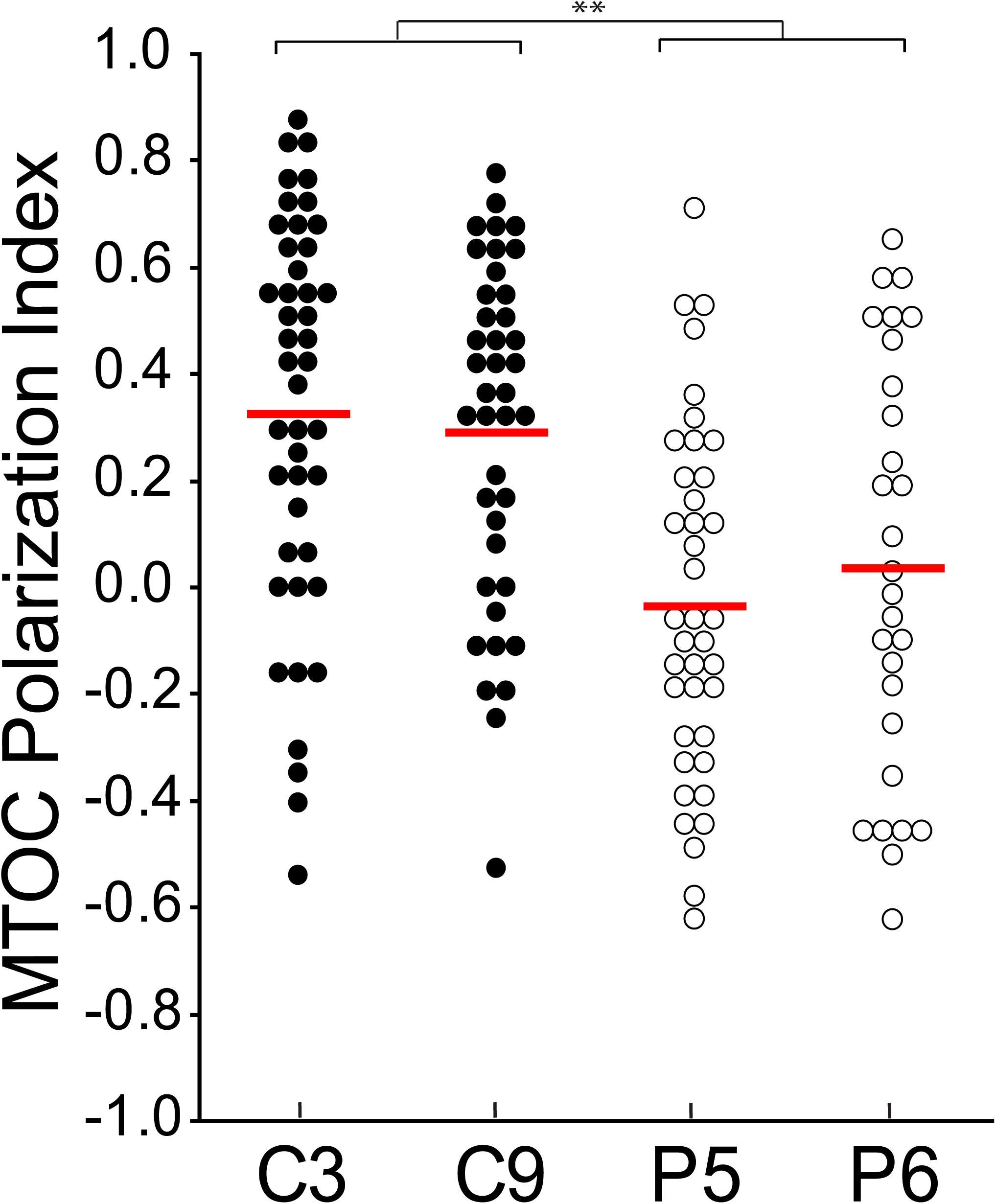
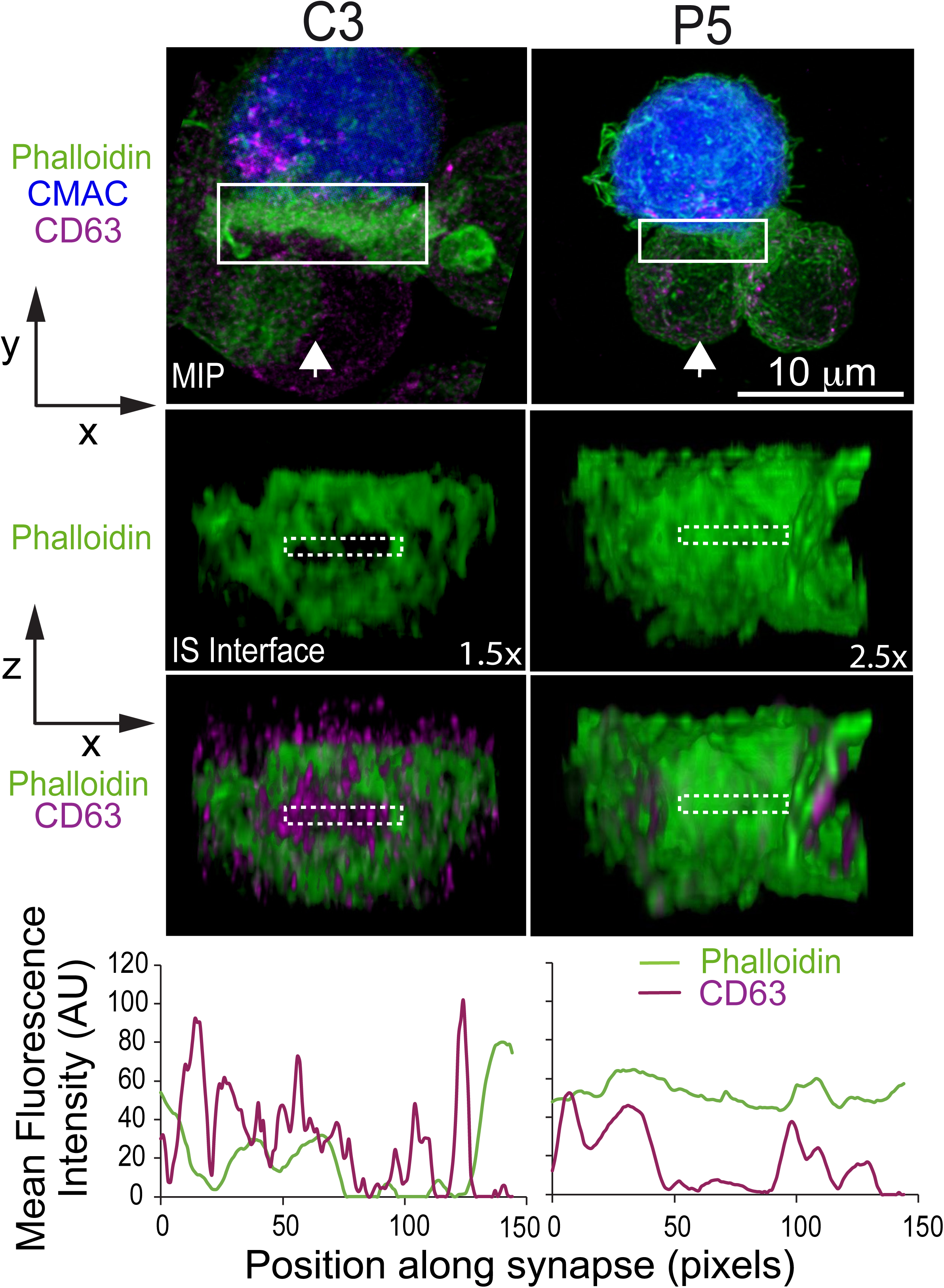
PKCδ regulates MTOC and MVB polarization. Panel A), C3 control and P5 PKCδ-interfered clones were challenged with CMAC-labelled SEE-pulsed Raji cells for 1 h, fixed, stained with anti-γ-tubulin AF546 to label MTOC and anti-CD63 AF647, to label MVB and imaged by confocal fluorescence microscopy. In the upper diagram, the distances (A, green; B, blue) used for the calculation of Pol. Index (A/B) are indicated. The red dot represents the cell center of mass (Cell^C^), whereas the yellow dot indicates the projection of MTOC or MVB center of mass (MTOC^c^ and MVB^c^, respectively) on the vector defined by the Cell^C^–synapse axis. Since the Cell^C^ position was taken as the origin to measure distances, those “A” distance values in the opposite direction to the synapse were taken as negative. Thus, polarization indexes range form +1 (fully polarized) to -1 (fully antipolarized). In the lower panels, MIP of the indicated, merged channels of representative images for both C3 and P5 forming synapses are included. The Raji cells and the Jurkat clones are labelled with discontinuous and continuous white lines, respectively. The superimposed white crosses label the MTOC^c^ or MVB^c^. The white arrow indicates MTOC position, whereas the yellow arrow labels the MVB. Panel B), MTOC Pol. Index was calculated as indicated in panel A for the indicated number of synaptic conjugates made by C3, C9 (control) and P5, P6 (PKCδ-interfered) clones, that were previously challenged for 1 h with SEE-pulsed Raji cells. Dot plot distribution and average Pol. Index (red horizontal line) are represented. The blue line indicates the cut-off value (Pol. Index = 0.25) used to calculate the percentage of synapses showing polarized MTOC. **, p ≤0.05. Panel C), C3 control and P5 PKCδ-interfered clones were challenged with CMAC-labelled SEE-pulsed Raji cells for 1 h, fixed, stained with phalloidin AF488 and anti-CD63 AF647 and imaged by confocal fluorescence microscopy. Upper panels: top views corresponding to the Maximal Intensity Projection (MIP) of the indicated, three merged channels, in a representative example. White arrows indicate the direction to visualize the face on views of the synapse (IS interface) enclosed by the ROIs (white rectangles) as shown in Suppl. Video 3. Middle panels: face on views of the IS. The enlarged ROIs from the upper panel (1.5x and 2.5x zoom, respectively) were used to generate (as shown in Suppl. Video 3) the IS interface merged images (phalloidin and CD63 channels from frame no. 43 of Suppl. Video 3). Lower diagrams: phalloidin and CD63 MFI vs position along the indicated, rectangular ROIs (discontinuous line) embed in the face on views from the lower panels. CMAC labelling of Raji cells in blue, phalloidin in green and CD63 in magenta. Scale bars, 10 µm.

**Fig. 2.**
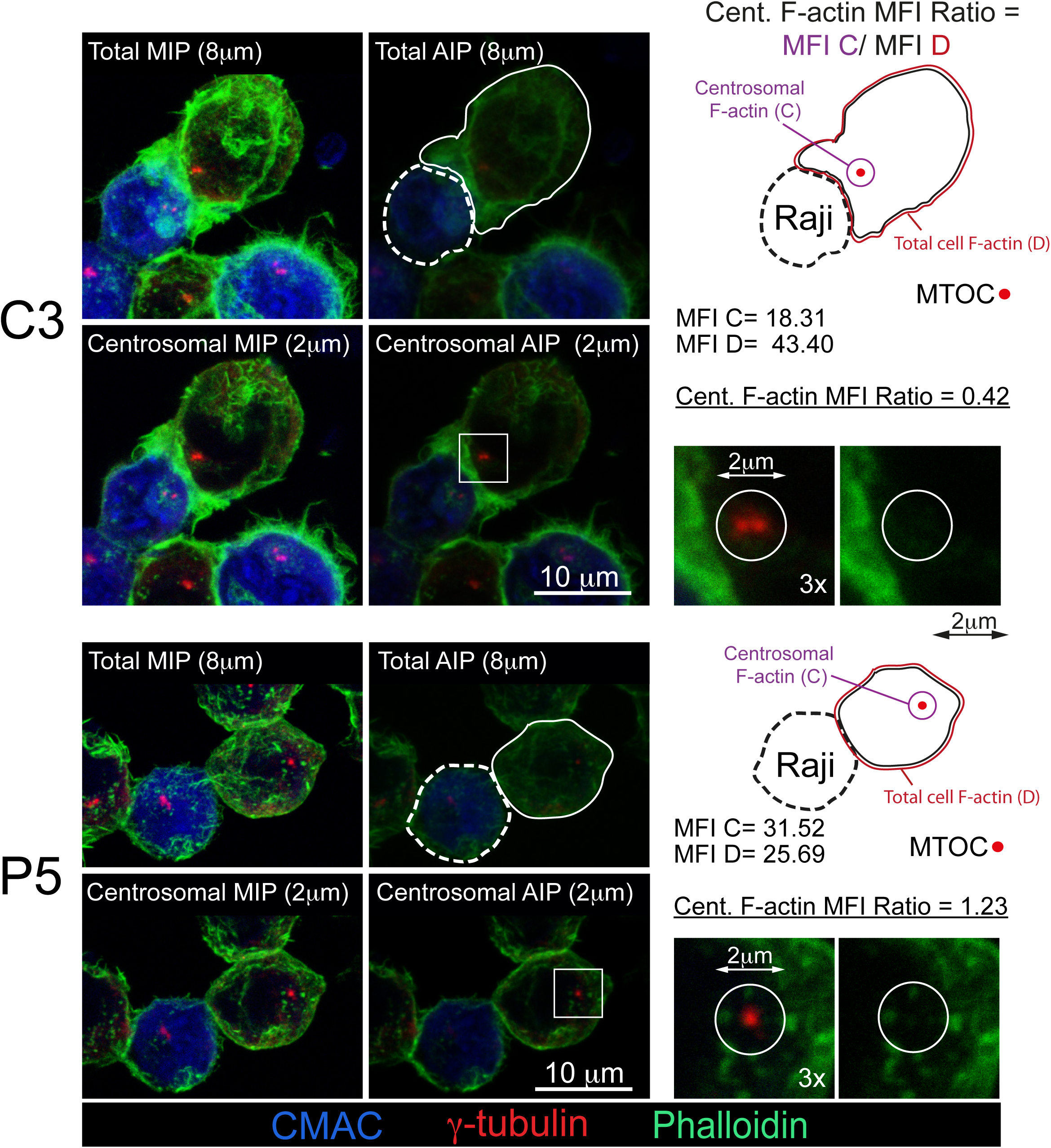
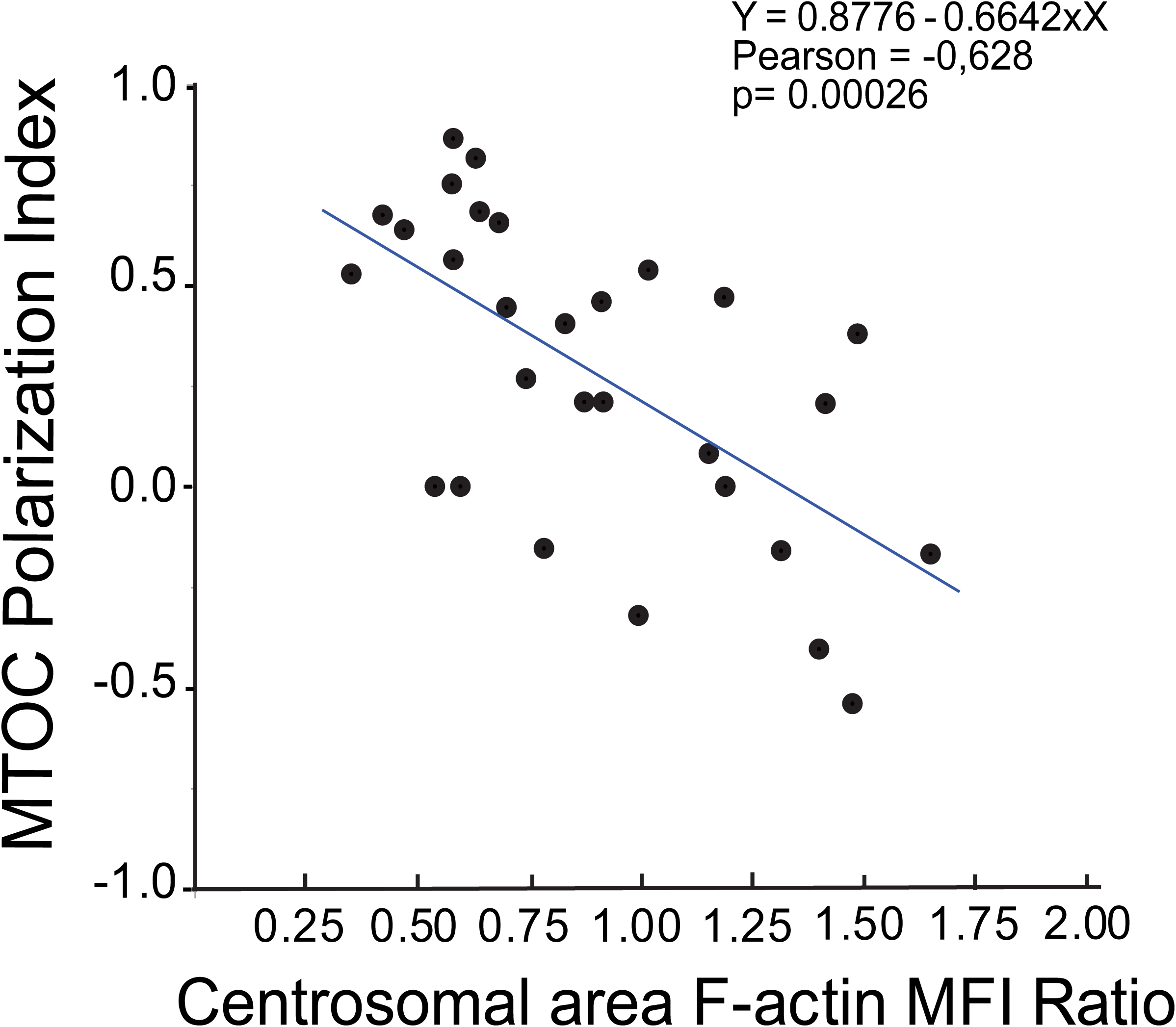
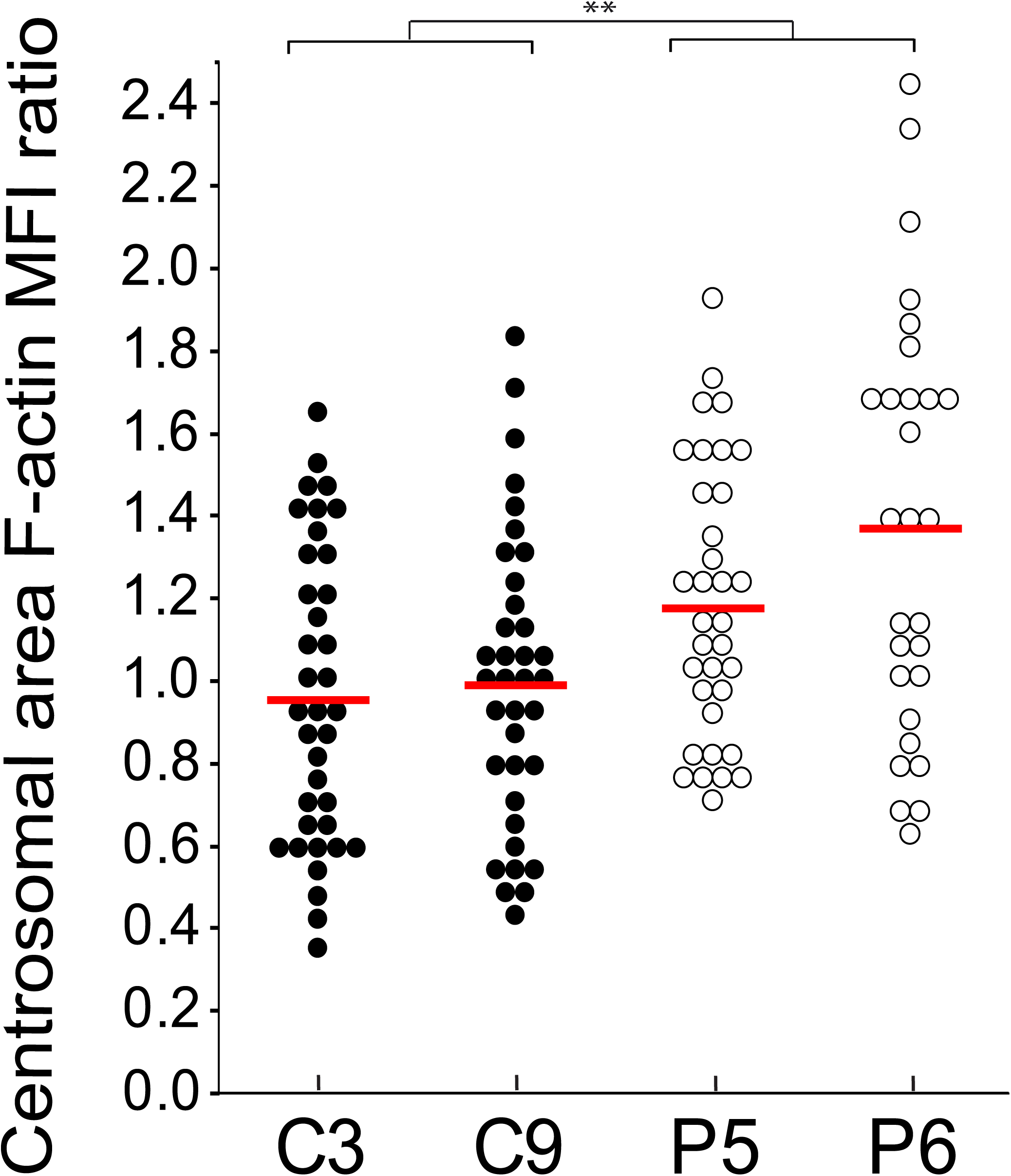
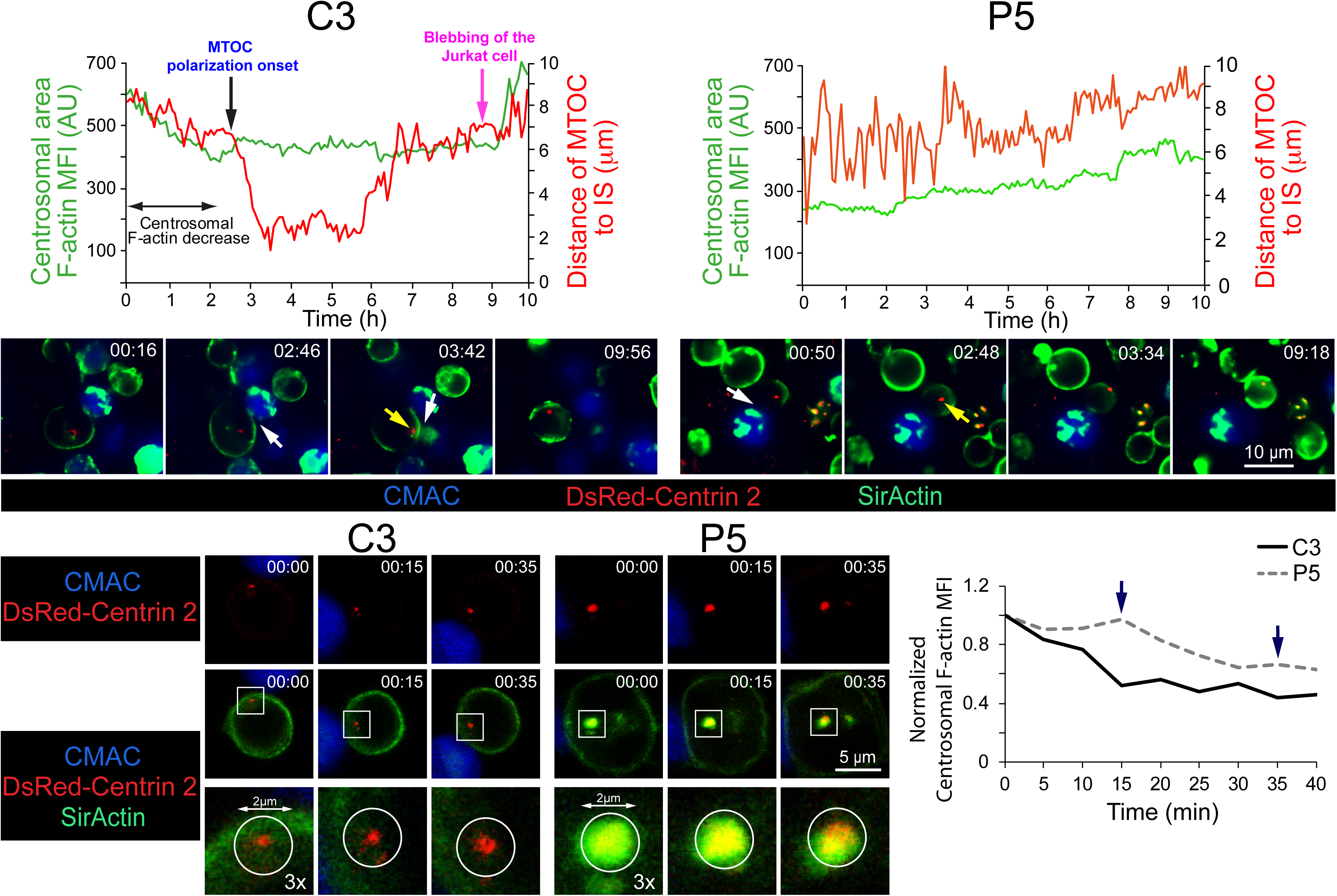
PKCδ regulates centrosomal area F-actin. Panel A), C3 control and P5 PKCδ-interfered clones were challenged with CMAC-labelled SEE-pulsed Raji cells for 1 h, fixed, stained with anti-γ-tubulin AF546 to label MTOC and phalloidin AF488 to label F-actin, and imaged by confocal fluorescence microscopy. Left, total (8 μm thick) or centrosomal (2 μm thick), maximal or average intensity projections (MIP and AIP, respectively) are shown as indicated. In the right side images, enlarged centrosomal areas (as defined in centrosomal AIPs, white squares) are shown containing the 2 μm-diameter centrosomal ROIs (white circles) used to calculate the centrosomal area F-actin MFI. In the right side diagrams, the F-actin MFI values in ROIs (“C” magenta ROI labels centrosomal area F-actin, whereas “D” red ROI labels total cell F-actin) used to calculate the centrosomal area F-actin MFI ratio are indicated. The Raji cells and the Jurkat clones are labelled with discontinuous and continuous lines, respectively. Panel B), C3 control clone was challenged with CMAC-labelled SEE-pulsed Raji cells for 1 h, fixed, stained with anti-γ-tubulin AF546 to label MTOC, and phalloidin AF488 to label F-actin and imaged by confocal fluorescence microscopy. Subsequently, centrosomal area F-actin MFI ratio and MTOC polarization index were calculated as indicated in Material and Methods, linear correlation analyses between these values corresponding to each individual cell was represented and Pearson’s correlation coefficient was calculated. Non-parametric Spearman’s correlation coefficient between these variables was ρ=-0.633 (p= 0.00017). Panel C), centrosomal area F-actin MFI ratio was calculated for synaptic conjugates made by C3, C9 (control) and P5, P6 (PKCδ-interfered) clones as indicated in panel A. Dot plot distribution and average centrosomal area F-actin MFI ratio (red horizontal line) are represented. **, p ≤0.05. Panel D). Upper diagrams, C3 control and P5 PKCδ-interfered clones expressing dsRed-Cent2 and pulsed with SirActin and verapamil were challenged with CMAC-labelled SEE-pulsed Raji cells bound to glass-bottom plates. Subsequently, synapses were imaged by wide-field, time-lapse microscopy and centrosomal area F-actin MFI (green line) and the distance of the MTOC to the IS (red line) were measured in each frame, as indicated in Materials and Methods, using a 2 μm-diameter floating ROI, centered at the MTOC^C^. Vertical black arrow labels the onset of MTOC polarization, whereas magenta arrow labels the blebbing of the Jurkat clone at late time points. Horizontal double-arrow line labels the time period during which centrosomal area F-actin decreased. Middle panels, some representative frames of the indicated, merged channels (CMAC, blue; dsRed-Cent2, red; SirActin, green) are included (see also Suppl. Video 4). White arrow labels the synapse, whereas yellow arrow labels the MTOC. Lower panels, C3 control and P5 PKCδ-interfered clones expressing dsRed-Cent2 and pulsed with SirActin and verapamil were challenged with CMAC-labelled SEE-pulsed Raji cells in suspension. Subsequently, synapses were imaged by confocal time-lapse microscopy and centrosomal area F-actin MFI was measured, using a 2 μm-diameter floating ROI centered at the MTOC^C^, as indicated in Materials and Methods. Left panels include some representative frames of the merged channels (CMAC, blue; dsRed-Cent2, red; SirActin, green, see also Suppl. Video 5), whereas the right diagram represents the kinetics of normalized centrosomal area F-actin MFI (referred to MFI at t=0). Dark arrows indicate the representative frames shown in the left panels. The slow decrease in centrosomal area F-actin MFI occurring in P5 clone, which was not evident in the epi-fluorescence experiment shown in the upper plot, is due to the increased fluorescence bleaching by the confocal illumination conditions. Data are representative of the results obtained in several experiments (*n* = 3).

## RESULTS

### PKCδ interference decreases both MTOC and MVB polarization towards the IS

We have previously shown, by using a well-established IS model [31] [35] [32], that PKCδ-interfered Jurkat clones challenged with antigen on APC secreted a lower amount of exosomes upon IS formation [14]. We also defined that the deficient polarization of MVB containing the exosome precursors towards the IS underlies the decreased exosome secretion [14]. In the context of secretory traffic at the IS, it has been published that not always MTOC polarization is necessary or sufficient for the transport or certain secretory vesicles to the IS [7], such as lytic granules, and for the cytotoxic hit delivery [36–38]. All these previous examples of segregation between MTOC movement and secretory granules traffic prompted us to simultaneously analyze both MTOC and MVB movement at the single cell level. We first analysed by time-lapse microscopy MTOC polarization, MVB convergence towards the MTOC and MVB polarization in Jurkat clones co-expressing dsRed-Cent2 and CFP-CD63, upon challenge with SEE-pulsed Raji cells, which is a well-established Th synapse model [31]. As shown in the representative Suppl. Video 1, MVB convergence towards the MTOC occurred almost simultaneously with MTOC migration towards the IS. We then quantitated MTOC and MVB polarization in fixed synapses formed by control or PKCδ-interfered Jurkat clones by fluorescence microscopy, as previously described [14] (Fig. 1A). As seen in Fig. 1B, C3 and C9 control clones substantiated significantly higher MTOC polarization indexes when compared to P5 and P6 PKCδ-interfered clones. We have previously compared MVB polarization indexes among these clones obtaining comparable results [14]. Remarkably, we found a strong linear correlation between MVB and MTOC polarization indexes (Pol. Index) for each clone (i.e. Pearson’s linear correlation coefficient 0.962 and 0.949 for C3 and P5 clones, respectively, Suppl. Fig S1), accordingly with the Suppl. Video 1. Furthermore, in all the analyzed synapses the MTOC^C^ was coincident or very proximal to the MVB^C^ (i.e. Suppl. Video 1, Fig. 1A), regardless of polarization. Although the MTOC and MVB did not efficiently polarize in P5 and P6 PKCδ–interfered cells (Fig. 1), their MVB still converged towards the MTOC (Fig. 1A, Suppl. Fig. S1). Thus, PKCδ appears to specifically regulate MTOC movement to the IS.

### PKCδ regulates F-actin reorganization at the IS and around the MTOC during MTOC polarization

Our previous results show that PKCδ regulates cortical F-actin reorganization at the IS, contributing to MVB polarization and exosome secretion [14], since PKCδ-interfered clones forming synapses exhibited quantitative alterations (duration and magnitude of F-actin reorganization), but also qualitative differences (absence of depletion of F-actin at the cIS) in F-actin reorganization at the IS when compared with control clones [14]. All these defects were rescued by ectopic, mouse PKCδ re-expression in PKCδ-interfered clones, as well as the defective MTOC/MVB polarization [14]. We aimed to explore the consequences of this defective cortical F-actin reorganization on MTOC/MVB polarization. To this end, we labelled C3 control and P5 PKCδ-interfered clones with phalloidin and anti-CD63, to visualize in the same cell F-actin and MVB, respectively, by confocal microscopy. We then analyzed the 3D distributions of F-actin and MVB and generated 2D projections of both distributions at the IS interface (Suppl. Video 3 and Fig. 1C, medium and lower panels). Next, to evaluate the relative position of CD63^+^ vesicles (MVB) with respect to F-actin architecture at the IS, the MFI of F-actin and CD63 along the IS interface (ROI labelled with discontinuous white-line rectangles in Fig. 1C) was measured (Fig. 1C, bottom diagrams). In C3 control clone we found an accumulation of MVB (peaks in the magenta MFI profile, lower panels) included in the F-actin-depleted area (represented by a valley in the green MFI profile) at the cIS (Fig. 1C), that includes the secretory domain [4]. In contrast, in P5 PKCδ-interfered clone, neither the F-actin-depleted area nor MVB were observed at the cIS (Fig. 1C). Thus, cortical F-actin reorganization at the IS is regulated by PKCδ, and this reorganization appears to control MVB polarization and hence MVB secretion. In addition, when we expressed dsRed-Cent2 in Jurkat clones to assess MTOC position in relation to the F-actin depleted area at the cIS, we found in C3 control clone the MTOC located at the edge of the cIS (Suppl. Fig. S2, yellow arrow in upper left panel) that corresponded to the described secretory domain next to the cSMAC [4], in accordance with the observed MVB position (Fig. 1C). However, in P5 PKCδ-interfered clone, the MTOC position was distal to the IS (Suppl. Fig. S2, yellow arrow in right panel). Thus, the spatial organization of F-actin at the IS is altered in PKCδ-interfered clones, and this may contribute to the deficient MTOC/MVB polarization, although at this stage we could not exclude that other non-cortical F-actin-dependent events may also contribute to the observed phenotype.

In this context, it has been demonstrated that F-actin depletion at a 2 μm –diameter centrosomal area appears to be crucial to allow MTOC polarization towards the IS in BCR-stimulated B lymphocytes [28]. However, to date no evidence of centrosomal area F-actin depletion has been reported during MTOC polarization in T lymphocytes. We thus analyzed centrosomal area F-actin in T lymphocytes in a control clone forming synapses. To perform these experiments we measured thresholded, F-actin MFI within a defined, spherical ROI (2 μm diameter) centered at the MTOC^c^, as previously described [28], and we divided this centrosomal area F-actin MFI by total cellular F-actin MFI, to normalize by cell area, shape and phalloidin staining variations among samples (Fig. 2A, upper panels). The possible correlation of the centrosomal area F-actin MFI ratio with the MTOC polarization index was analysed in C3 control (Fig. 2B) and P5 PKCδ-interfered (not shown) clones. A negative correlation was observed in the C3 control clone (Spearman’s Rho Coefficient = −0.632; p=0.00017 and Pearson’s Coefficient = −0.628; p=0.00026), but no correlation was observed (Spearman’s Rho Coefficient = −0.13; p=0.44 and Pearson’s Coefficient = −0.2; p=0.22) in the P5 PKCδ-interfered clone. Moreover, when we compared control and PKCδ-interfered clones forming synapses in end point experiments, we found a statistically significant higher centrosomal area F-actin MFI ratio in PKCδ-interfered clones (Fig. 2A and 2C). Thus, PKCδ-interfered clones forming synapses exhibited higher levels of centrosomal area F-actin.

The asynchronous character of synapse formation may also lead to asynchronous MTOC polarization [39] [35] [14], that may in turn skew the results from end point, fixed synapses. To circumvent this caveat, we first analyzed by epifluorescence and subsequent image deconvolution, the centrosomal area F-actin reorganization upon synapse formation by time-lapse analyses at the single, living-cell level. To correlate centrosomal area F-actin with MTOC polarization we performed these experiments with clones expressing dsRed-Cent2 [39], loaded with SirActin and verapamil [40] and subsequently challenged with SEE-pulsed Raji cells. As seen in Fig. 2D and Suppl. Video 4, upon the synaptic contact of C3 control clone (white arrow in Suppl. Video 4), concomitantly with the initial increase in cortical F-actin MFI at the IS [14] (Suppl. Video 4 and Fig. 2D, white arrow in the middle panel), a progressive decrease of centrosomal area F-actin MFI (labelled by a double arrow line in Fig. 2D, upper left panel) was observed, which represented centrosomal area F-actin dismantling. Neither centrosomal area F-actin dismantling nor MTOC polarization occurred in control clones not forming synaptic conjugates (not shown). Remarkably, the early decrease in centrosomal area F-actin MFI occurred before the onset of the rapid polarization of MTOC towards the IS (black vertical arrow in Fig. 2D, upper-left panel). This was consistent with the hypothesized role of centrosomal area F-actin dismantling in MTOC polarization. At late time points (magenta arrow in Fig. 2D and Suppl. Video 4), plasma membrane blebbing (an early feature of T lymphocyte AICD) occurred in the C3 control clone forming synapse, as previously shown [14] [15] [17]. Centrosomal area F-actin dismantling did not occur in the P5 PKCδ-interfered clone, since centrosomal area F-actin MFI was maintained (Fig. 2D, upper-right panel), concomitantly with the absence of MTOC polarization (yellow arrow in Supp. Video 4, lower panels, and right-middle panel in Fig. 2D) and no AICD occurred (Suppl. Video 4). End point analyses of centrosomal area F-actin and MTOC polarization in dsRed-Cent2-expressing clones rendered similar results to those obtained by labelling MTOC with anti-γ-tubulin (compare Suppl. Fig. S3 with Fig. 2A and 2C). In addition, to enhance the spatial resolution we performed shorter time-lapse analyses but using confocal microscopy. As seen in Suppl. Video 5, the P5 PKCδ-interfered clone forming synapses exhibited higher levels of centrosomal area F-actin than the C3 control clone (Fig. 2D, lower right panel). Centrosomal area F-actin partially overlapped with dsRed-Cent2 in the P5 PKCδ-interfered clone, but not in the C3 control clone (Fig. 2D, lower left panel). The high centrosomal area F-actin green fluorescence partially overlapped the dsRed-Cent2 red fluorescence at MTOC, rendering an evident yellow dot in the P5 clone (Fig. 2D, lower left panel). Thus, centrosomal area F-actin dismantling correlated with MTOC polarization to the IS and, since PKCδ-interfered clones exhibited sustained levels of centrosomal area F-actin, PKCδ appeared to negatively regulate this process.

### FMNL1 colocalizes with F-actin at the IS but not around the MTOC

FMNL1 is required for MTOC polarization in T lymphocytes and this event appears to be independent of Arp2/3-mediated, cortical F-actin reorganization at the IS [23]. In PKCδ-interfered clones we have observed that the spatial F-actin organization at the IS is affected, but also the centrosomal area F-actin reorganization (see above). Thus, it is conceivable that PKCδ may exert its regulatory role on MTOC polarization via regulation of FMNL1 function acting separately and/or coordinately in any of these subcellular localizations. FMNL1 is located at the IS and around the MTOC [23] and PKCδ is located around the MTOC [14]. As a first approach to understand the molecular bases of PKCδ effect on the different, subcellular F-actin networks, we studied subcellular localization of FMNL1 in relation to F-actin cytoskeleton at both the IS and centrosomal area. As shown in Figs. 3A and 3B, FMNL1 and F-actin partially colocalized at the IS, similarly in C3 control and P5 PKCδ-interfered clones (Fig. 3B). In addition, no colocalization of FMNL1 and F-actin was observed around the MTOC either in the C3 or the P5 clone (Fig. 3B and Suppl. Fig. S4).

**Fig. 3.**
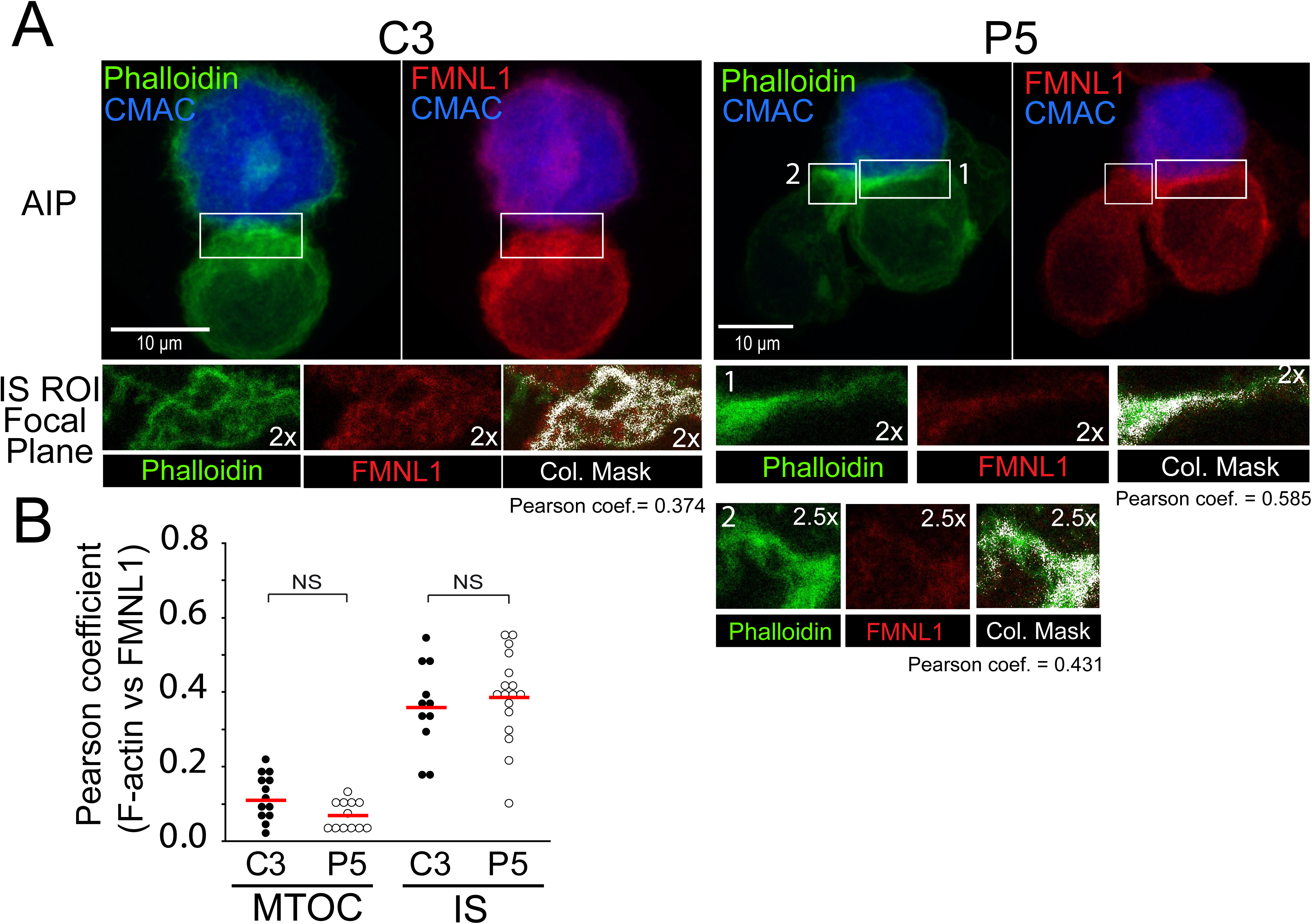
Subcellular localization of FMNL1. C3 control (left) and P5 PKCδ-interfered clones (right) were challenged with CMAC-labelled SEE-pulsed Raji cells for 1 h, fixed, stained with anti-FMNL1 AF546 and phalloidin AF488 to label F-actin, and imaged by confocal fluorescence microscopy. Panel A) In the upper panels, representative average intensity projections (AIP) of the indicated, merged channels of synaptic conjugates made by C3 (left panels) and P5 (right panels) clones are shown. White rectangles enclosed the IS ROIs used for the colocalization analyses of the indicated channels in the optical sections represented in the lower row of panels. In the right side of this row, colocalization masks corresponding to merged, phalloidin and FMNL1 channels, are represented in white. Pearson’s coefficients corresponding to each colocalization analysis are indicated below each colocalization mask panel. CMAC labelling of Raji cells in blue, phalloidin in green and FMNL1 in red. Panel B) same as panel A, but colocalization analyses were performed for both centrosomal, MTOC ROI (see also Suppl. Fig. S4) and IS ROI in both C3 control and P5 PKCδ- interfered clones. Dot plot distribution represents the Pearson’s coefficients corresponding to each colocalization analysis and to each clone. Data are representative of the results obtained in several experiments (*n* = 3). NS, not significant.

### PKCδ induces FMNL1 phosphorylation

Since no differences in the subcellular localization of FMNL1 with respect to F-actin between control and PKCδ-interfered clones were observed, and all the clones exhibited similar levels of FMNL1 and Dia1 (Fig. 4, upper panel and Suppl. Fig. S5), we aimed to search for potential post-translational modifications in FMNL1 and Dia1, such as phosphorylation, which may underlie the observed phenotype. FMNL2 is phosphorylated by PKCα and, to a lower extent, by PKCδ at S1072 [41], reversing its autoinhibition by the C-terminal, DAD auto-inhibitory domain and enhancing F-actin assembly, β1-integrin endocytosis, and invasive motility [41]. In FMNL1β, S1086 is surrounded by a sequence displaying high homology to the one surrounding S1072 of FMNL2, whereas the homology of FMNL1α and FMNL1γ with the C-terminus of FMNL2 was much lower [42] [41] (Suppl. Fig. S6). In addition, the three FMNL1 isoforms share identical sequence from amino acid residue 1 to 1070, and diverge in the C-terminal region, which includes the DAD auto-inhibitory domain (Suppl. Fig. S6). Thus, PKCδ may regulate F-actin reorganization by controlling FMNL1β S1086 phosphorylation, as certain PKC isoforms regulate FMNL2 activity [41]. We immunoprecipitated FMNL1 from C3 control and P5 PKCδ-interfered clones, untreated or treated with PKC activator PMA or anti-TCR, and analysed FMNL1 phosphorylation in the immunoprecipitates by WB with anti-Phospho-(Ser) PKC substrate antibody [41]; the results were normalised by FMNL1 levels in the immunoprecipitates. As shown in Fig. 4A, both PMA and anti-TCR stimulation induced FMNL1 phosphorylation in the C3 control clone to a higher extent than in the P5 PKCδ-interfered clone. Moreover, expression of an interference-resistant, GFP-PKCδ in the P5 PKCδ-interfered clone restored the phosphorylation of FMNL1 to the levels found in the C3 control clone (Fig. 4B). In contrast, when we analysed in parallel Dia1 phosphorylation using the same anti-Phospho-(Ser) PKC substrate antibody, we could not detect any phosphorylation of Dia1 induced by PMA in the Dia1 immunoprecipitates (Suppl. Fig. S5). Thus, PKC isoforms activated by PMA (including PKCδ) appear to regulate FMNL1, but not Dia1, phosphorylation. Since expression of an interference-resistant GFP-PKCδ restored FMNL1 phosphorylation (Fig. 4B) and MTOC polarization [14], all these data indicate that PKCδ regulates phosphorylation of FMNL1 and probably its function during MTOC polarization.

**Fig. 4.**
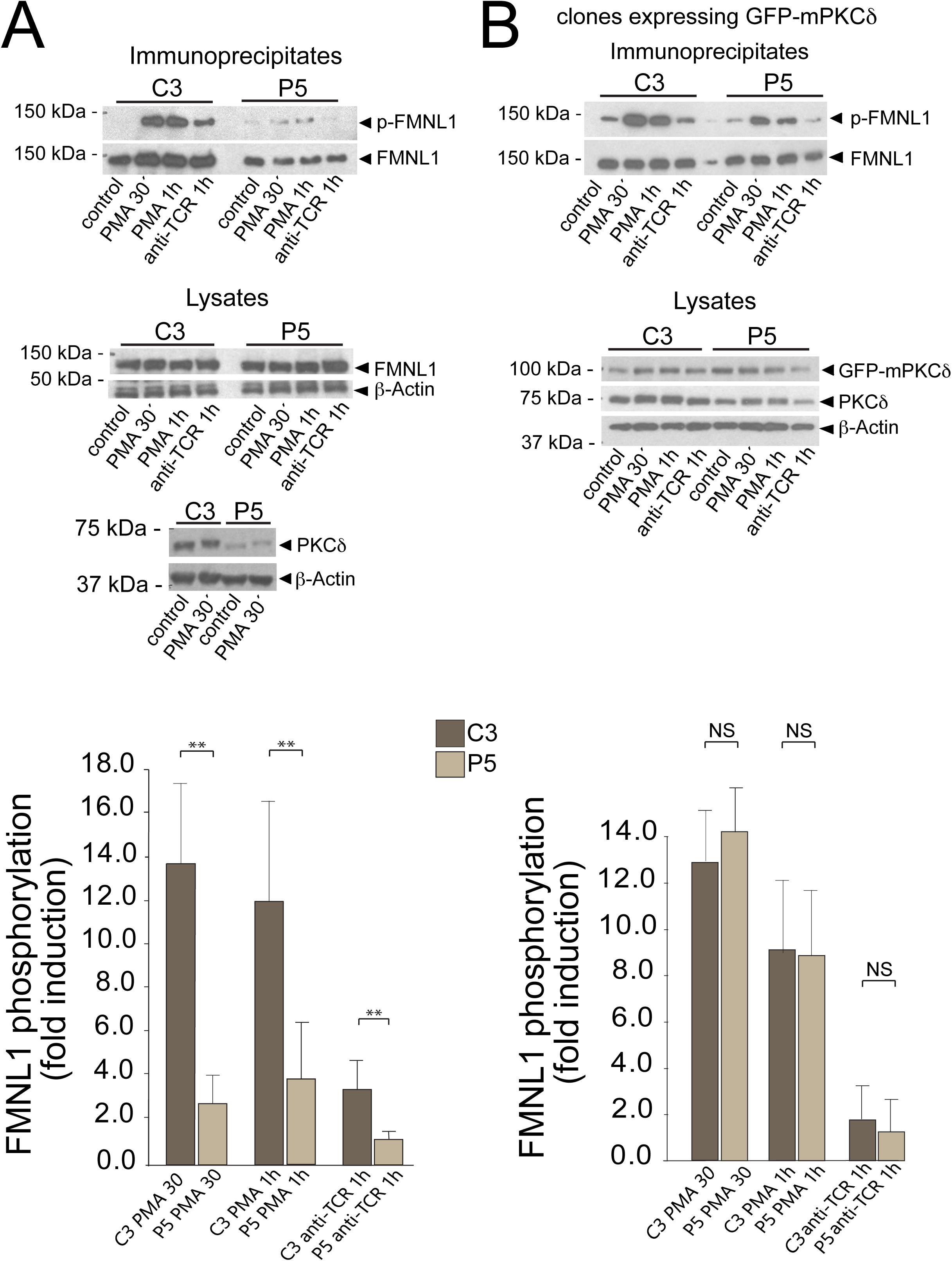
PKCδ regulates FMNL1 phosphorylation. Panel A), C3 control and P5 PKCδ-interfered clones were untreated or stimulated with either PMA or plastic-bound anti-TCR and subsequently lysed. WB of anti-FMNL1 immunoprecipitates (IPs) (upper panel) and cell lysates (lower panel) were sequentially probed with anti-Phospho-(Ser) PKC substrate and anti-FMNL1, to normalize Phospho-(Ser) PKC substrate signal for FMNL1 content in the IPs. In addition, cell lysates were probed with anti-FMNL1, β-actin and PKCδ (to check PKCδ interference). In the lower bar graph, the fold induction of FMNL1 phosphorylation, normalized to FMNL1 levels, was evaluated by WB quantification of several experiments similar to that described in the upper panel. Data are means plus SD of the results obtained in several (n = 5) experiments. **, p ≤0.05. Panel B) C3 control and P5 PKCδ-interfered clones were transfected with an interference resistant, GFP-mPKCδ expression plasmid. Subsequently, cells were stimulated as in panel A, and the anti-FMNL1 immunoprecipitates analysed by WB with anti-Phospho-(Ser) PKC substrate and anti-FMNL1, to normalize Phospho-(Ser) PKC signal for FMNL1 content in the IPs. In addition, the cell lysates were probed with anti-FMNL1, anti-β-actin and anti-panPKCδ, to check for both PKCδ interference and GFP-mPKCδ expression. In the lower bar graph, the fold induction of FMNL1 phosphorylation in cells expressing GFP-mPKCδ, normalized to FMNL1 levels, was evaluated by WB quantification of several experiments similar to that described in the upper panel. Data are means plus SD of the results obtained in several (n = 5) experiments. NS, not significant; **, p≤0.05.

### FMNL1β phosphorylation and MTOC polarization

The previous results suggest that PKCδ-dependent phosphorylation of FMNL1 may regulate FMNL1 function on F-actin reorganization at the IS and hence affect MTOC polarization. Is has been shown that FMNL1 interference impedes MTOC polarization [23]. Jurkat cells contain three FMNL1 isoforms (α, β and γ) [30] and the antibody we used for IP recognizes all these isoforms. Thus, any of these isoforms may be phosphorylated and/or involved in F-actin reorganization and hence MTOC polarization. To investigate the contribution of FMNL1 isoforms in MTOC polarization, we transfected the C3 control and P5 PKCδ-interfered clones with a bi-cistronic YFP expression plasmid interfering all FMNL1 isoforms (shFMNL1-HA-YFP) [30]. We then challenged the clones with SEE-pulsed Raji cells and analysed MTOC polarization in non-transfected (NT, YFP^-^) cells and FMNL1-interfered (YFP^+^) cells. As seen in Fig. 5A, the transient interference of all FMNL1 isoforms decreased the efficiency of MTOC polarization with respect to non-transfected C3 control cells forming synapses. However, it is noteworthy that FMNL1 interference did not further decrease MTOC polarization in the P5 PKCδ-interfered clone (Fig. 5B), which is compatible with PKCδ and FMNL1 being in the same regulatory pathway. We then transfected the C3 control clone with either a plasmid interfering all FMNL1 isoforms (shFMNL1-HA-YFP) or a plasmid interfering all FMNL1 isoforms and expressing interference-resistant FMNL1β (shFMNL1-HA-YFP-FMNL1β). YFP-FMNL1β expression rescued the defective MTOC polarization occurring in FMNL1-interfered cells (Fig. 5A) but it was not able to decrease the high levels of the centrosomal area F-actin to control levels (Fig. 5C). This suggests that, at least in conditions where all the FMNL1 isoforms, excluding FMNL1β, are interfered, centrosomal area F-actin decrease does not appear to be an absolute requirement for MTOC polarization. In addition, expression of interference-resistant FMNL1α (shFMNL1-HA-YFP-FMNL1α) or FMNL1γ (shFMNL1-HA-YFP-FMNL1γ) did not rescue the MTOC polarization to control levels (Suppl. Fig. S7), supporting a specific role of FMNL1β in MTOC polarization. Moreover, expression of FMNL1α or FMNL1γ was unable to rescue the centrosomal area F-actin to control levels (Suppl. Fig. S8), as it occurred with FMNL1β.

**Fig. 5.**
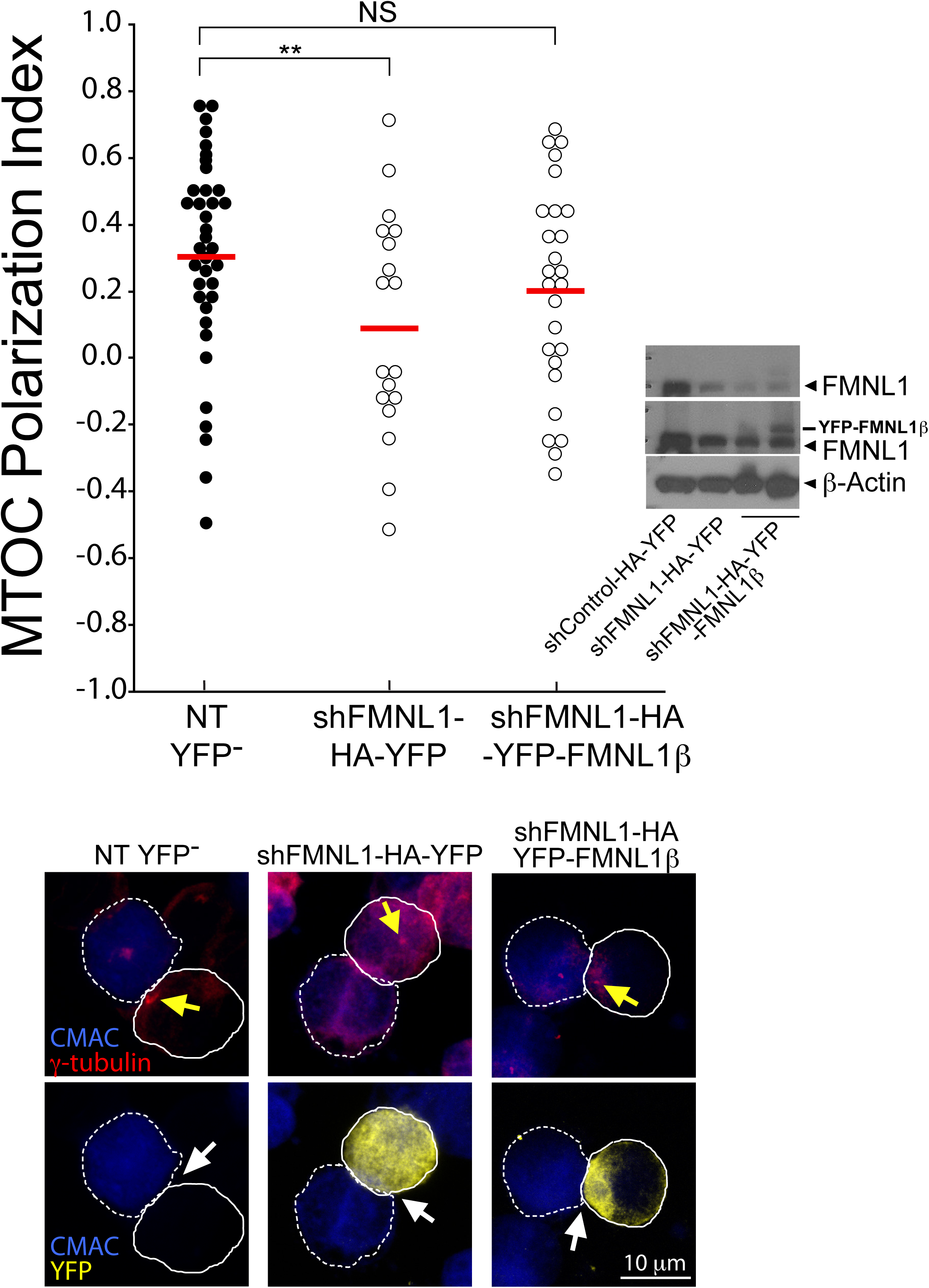
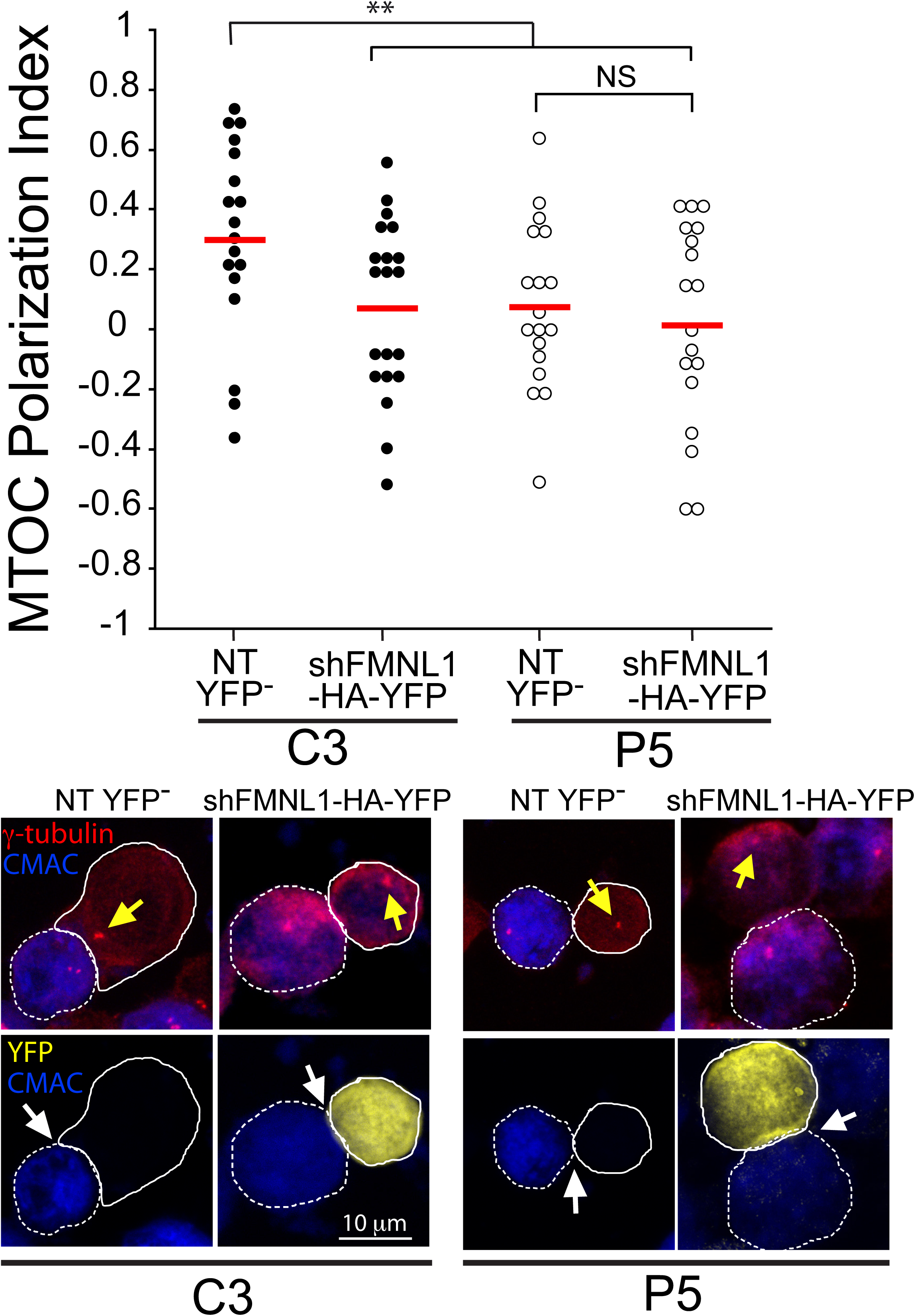
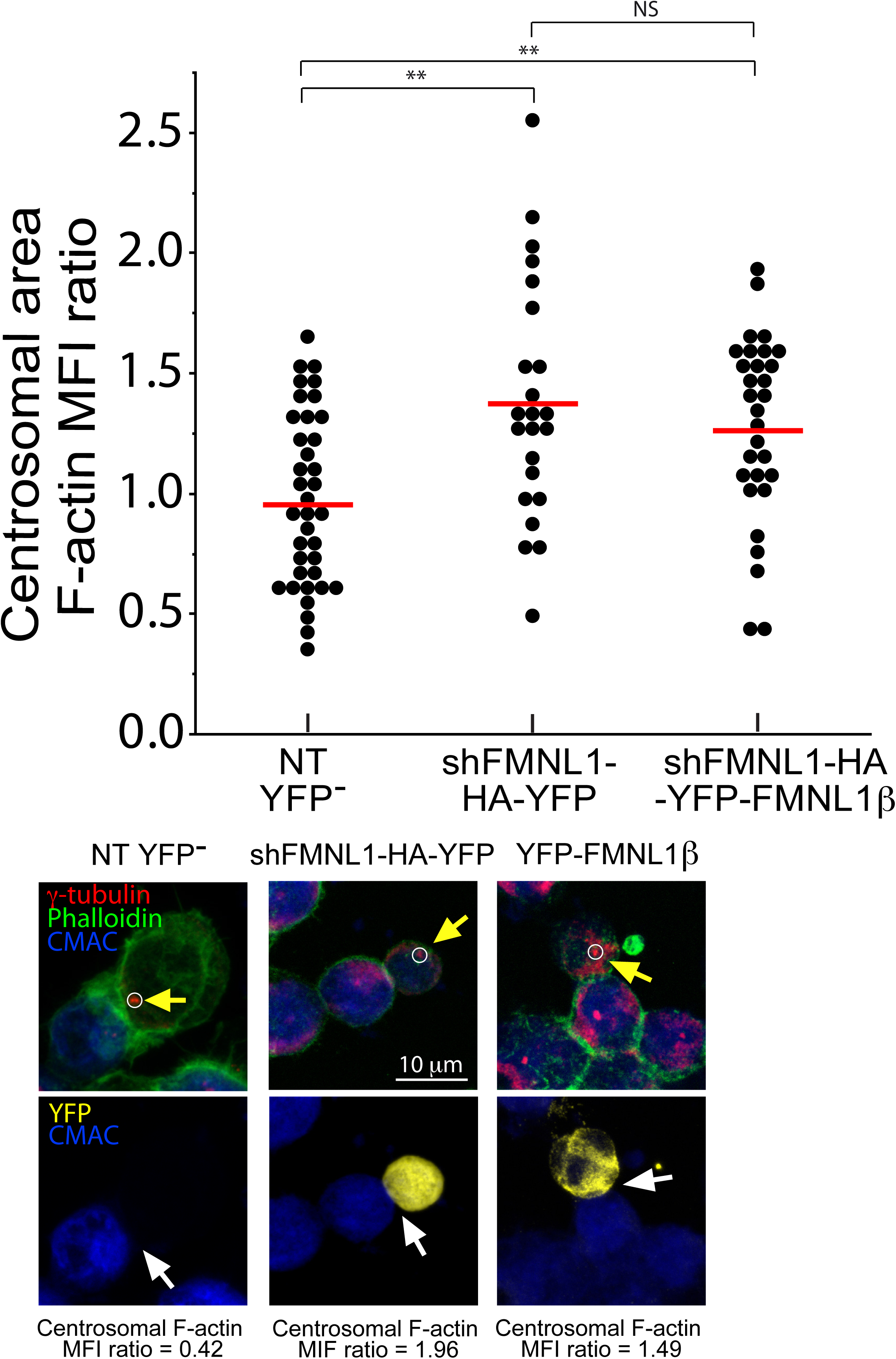
FMNL1 interference affects both centrosomal area F-actin and MTOC polarization. FMNL1β regulates MTOC polarization. C3 control and P5 PKCδ- interfered clones were transfected with either control (shControl-HA-YFP), FMNL1 interfering (shFMNL1-HA-YFP), or FMNL1-interfering, YFP-FMNL1β expressing vector (shFMNL1-HA-YFP-FMNL1β). Subsequently, the transfected clones were challenged with CMAC-labelled SEE-pulsed Raji cells for 1 h, fixed, stained with anti-γ-tubulin AF546 (red) and phalloidin AF647 (green) to label F-actin, and imaged by confocal fluorescence microscopy. Yellow channel fluorescence identifies the transfected cells. Panel A) upper diagram, MTOC Pol. Index was calculated as indicated above, for the indicated number of synaptic conjugates made by C3 control clone, transfected or not (NT YFP^-^ cells). Dot plot distribution and average Pol. Index (red horizontal line) are represented. In the inset, WB of cell lysates from the different groups of cells used was developed with anti-FMNL1 (two different expositions) and anti-β−Actin to check for both FMNL1 interference and HA-YFP-FMNL1β expression. Lower panels, representative synapses made by C3 clone, transfected or not (NT YFP^-^). This group includes all non-transfected cells from both shFMNL1-HA-YFP and shFMNL1-HA-YFP-FMNL1β transfections as internal controls. AIPs of the indicated, merged channels for C3 clone forming synapses are represented. Raji cells and Jurkat clones are labelled with discontinuous and continuous white lines, respectively. Panel B), upper diagram, MTOC Pol. Index was calculated as indicated in Fig. 1, panels A and B, for the synaptic conjugates made by C3, (control) and P5 (PKCδ-interfered) clones, transfected or not (NT YFP^-^). Dot plot distribution and average Pol. Index (red horizontal line) are represented.. Lower panels, representative synapses made by the different clones, transfected or not (NT YFP^-^). AIPs of the indicated, merged channels for both C3 and P5 clones forming synapses are represented. White arrows indicate the synaptic area, whereas yellow arrows indicate the MTOC position. Panel C), upper diagram, centrosomal area F-actin MFI ratio was calculated as indicated in Fig. 2 for the indicated number of synaptic conjugates made by C3 control clone, transfected or not (NT YFP^-^). The mean centrosomal area F-actin MFI ratio (red horizontal line) for each condition is represented. Lower panels, representative synapses made by C3 clone, transfected or not (NT YFP^-^). AIPs of the indicated, merged channels for C3 clone forming synapses are represented. The centrosomal area F-actin MFI ratio value is indicated for each condition. The MTOC is labeled with yellow arrow and the synapses with white arrows. The white circle enclosing the MTOC labels the ROI used to calculate the centrosomal F-actin MFI. NS, not significant. **, p ≤0.05.

To explore the contribution of cortical actin reorganization to MTOC polarization in these conditions, F-actin architecture at the cIS was analysed. Depletion of F-actin at cIS in FMNL1-interfered cells was lower than in non-transfected cells (Suppl. Fig. S9), as occurs in PKCδ-interfered cells [14], and FMNL1β re-expression rescued the F-actin depletion at the cIS (Suppl. Fig. S9). Thus, in these cells, FMNL1β acting on cortical F-actin appears to be sufficient for MTOC polarization, despite of the high levels of centrosomal area F-actin.

FMNL1β is a strong candidate to be regulated by PKCδ-mediated phosphorylation (see above). To test this hypothesis, we first analysed unstimulated or PMA-stimulated, HA-YFP-FMNL1β-transfected C3 control cells by immunofluorescence with the anti-Phospho-(Ser) PKC substrate antibody. The low transfection efficiency of the FMNL1 constructions (less than 10% of cells expressed the FMNL1 isoforms) made unfeasible the quantitative immunoprecipitation of these isoforms. Thus, strictly controlled, single-cell image analyses of Phospho-(Ser) PKC-substrate fluorescence signal were required to address FMNL1 isoforms phosphorylation. PMA increased the Phospho- (Ser) PKC-substrate MFI signal in YFP-FMNL1β^+^ cells with respect to YFP^-^ cells (Suppl. Fig. S10A), but this increase did not occur in HA-YFP-FMNL1α, HA-YFP-FMNL1γ or HA-YFP-FMNL1ΔFH2 mutant (a deletion mutant lacking the C-terminal region including the S1086)-expressing cells (Suppl. Fig. S10B). The same analysis was performed for unstimulated or PMA-stimulated, HA-YFP-FMNL1β-transfected C3 control and P5 PKCδ-interfered cells and the anti-Phospho-(Ser) PKC-substrate signal in PMA-stimulated C3 control cells was significantly higher than in PMA-stimulated P5 PKCδ-interfered cells (Fig. 6A). Moreover, a linear correlation was observed between HA-YFP-FMNL1β expression and anti-Phospho-(Ser) PKC-substrate labelling in PMA-stimulated HA-YFP-FMNL1β-expressing, C3 control cells, and this correlation was not observed in PMA-stimulated, HA-YFP-FMNL1β- expressing P5 PKCδ-interfered cells (Suppl. Fig. S10C). In addition, colocalization of HA-YFP-FMNL1β and anti-Phospho-(Ser) PKC substrate signals was observed in PMA-stimulated HA-YFP-FMNL1β-expressing C3 control cells (Pearson coefficient >0.90, not shown). All these results, together with the fact that FMNL1 has no kinase activity, support a deficient specific phosphorylation of HA-YFP-FMNL1β in PMA-stimulated, PKCδ-interfered cells. In addition, the increase in anti-Phospho-(Ser) PKC substrate signal triggered by immune synapse formation in HA-YFP-FMNL1β- expressing, C3 control cells was higher than in YFP^-^, C3 control cells (Suppl. Fig. S10D). The anti-Phospho-(Ser) PKC-substrate signal in HA-YFP-FMNL1β- expressing, P5 PKCδ-interfered cells forming synapses was lower than in HA-YFP-FMNL1β-expressing, C3 control cells forming synapses (Fig. 6B). To support that part of the anti-Phospho-(Ser) PKC-substrate signal indeed corresponded to the phosphorylation of HA-YFP-FMNL1β, we performed a colocalization analysis and found that HA-YFP-FMNL1β, but not HA-YFP-FMNL1α, partially colocalized with the anti-Phospho-(Ser) PKC substrate signal at the IS (Fig. 6C). As an internal colocalization positive control we determined the Pearson colocalization coefficient of HA-YFP-FMNL1β versus anti-FMNL1 (>0.80, not shown). Thus, PKCδ interference appeared to specifically inhibit FMNL1β phosphorylation.

**Fig. 6.**
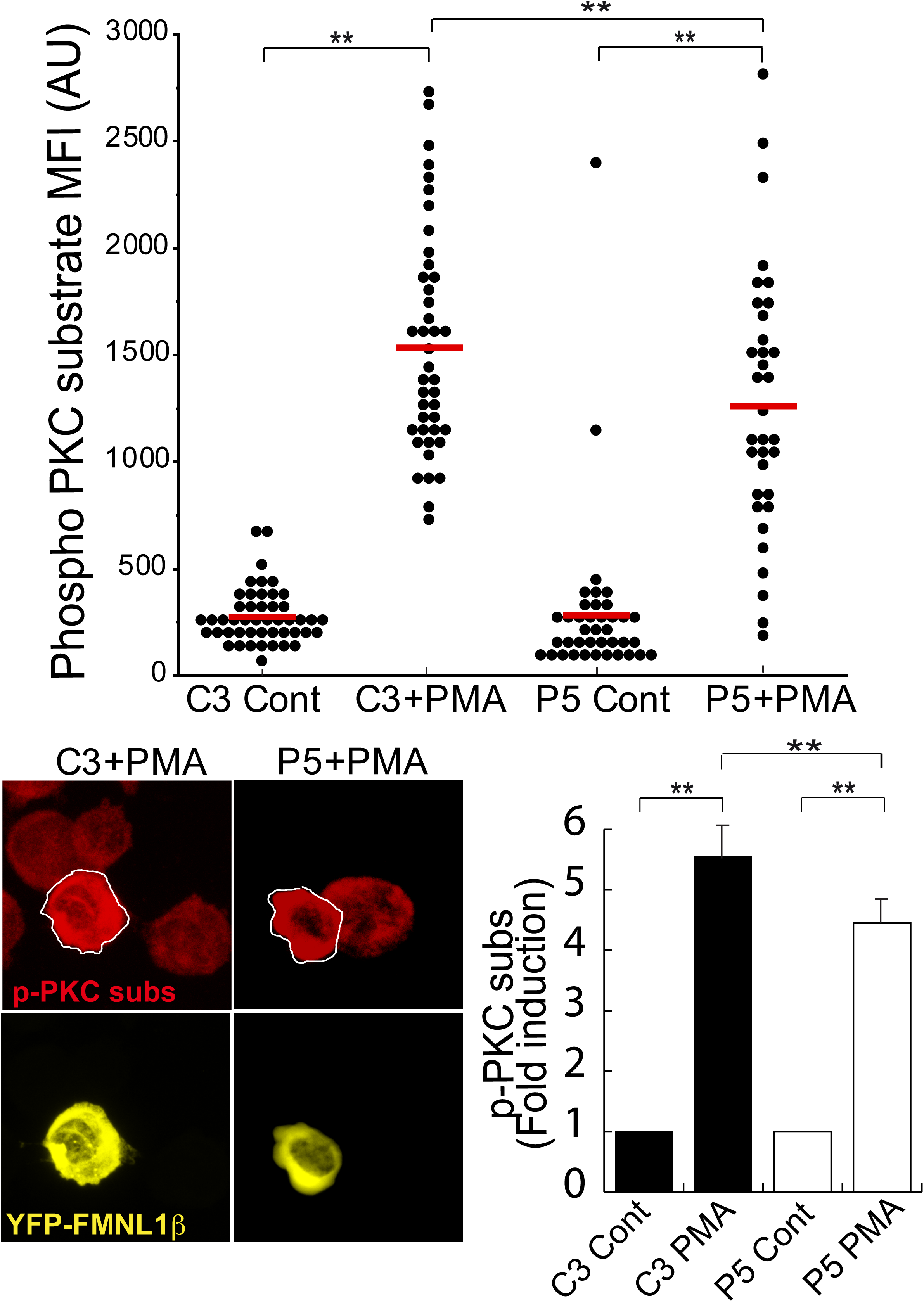
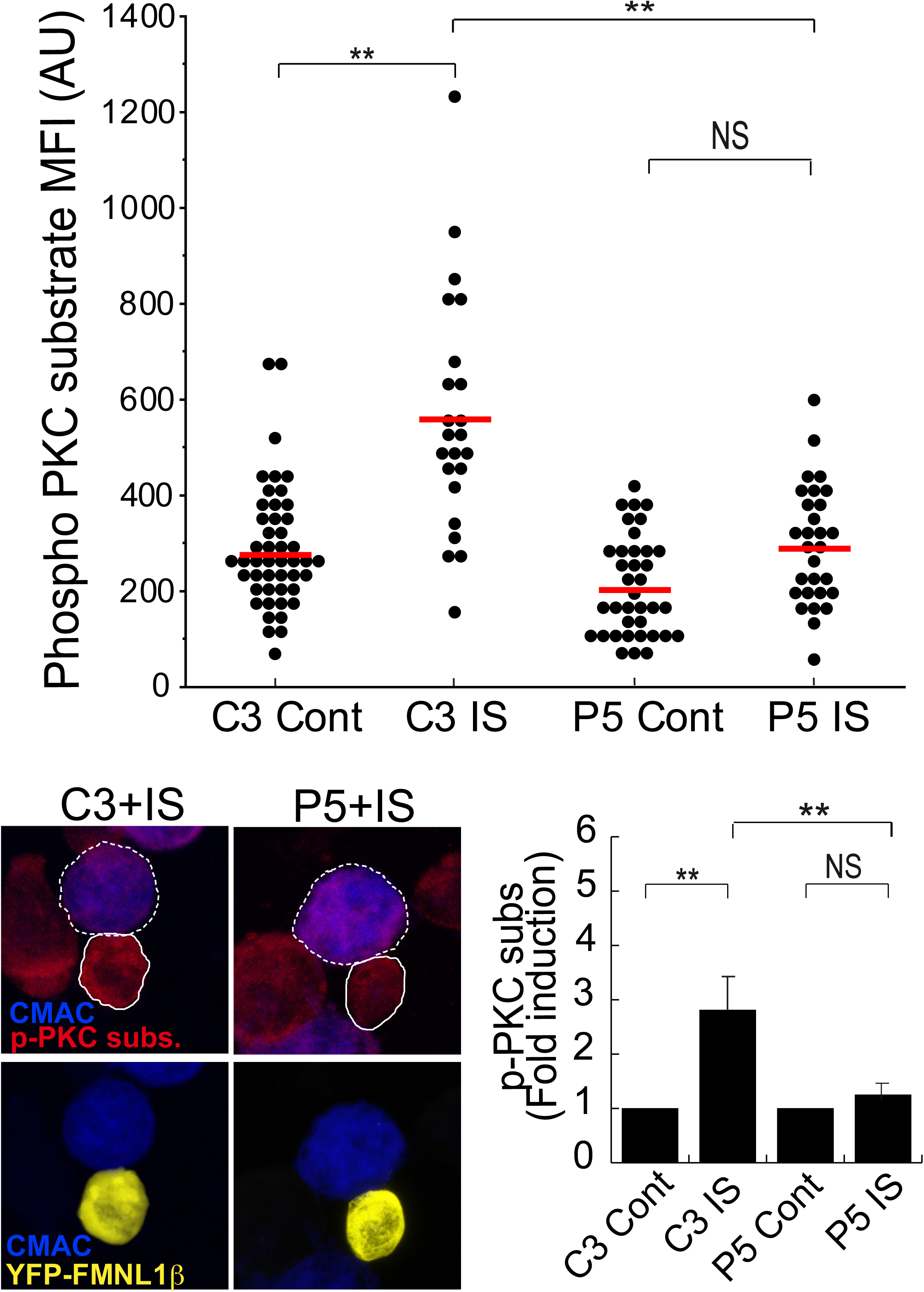
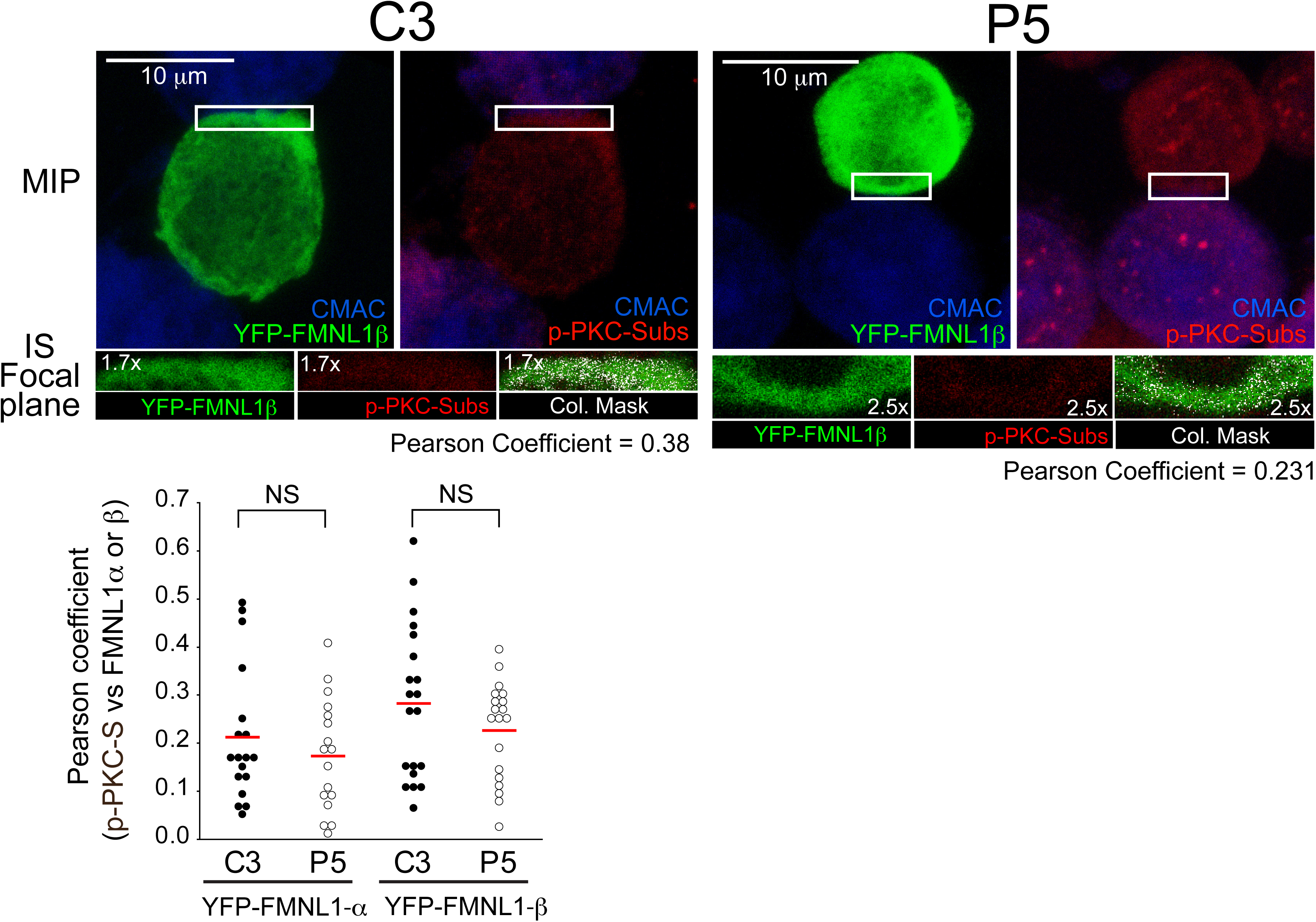
PKCδ regulates FMNL1β phosphorylation. C3 control and P5 PKCδ-interfered clones were transfected with an FMNL1-interfering, HA-YFP-FMNL1β- expressing expressing vector (shFMNL1-HA-YFP-FMNL1β). Subsequently, the transfected clones were either untreated (Cont), or stimulated with PMA (30 min), or challenged with CMAC-labelled SEE-pulsed Raji cells for 1 h, fixed, stained with anti-Phospho-(Ser) PKC substrate AF647 (magenta) and imaged by confocal fluorescence microscopy. Yellow channel fluorescence identifies the HA-YFP-FMNL1β−expressing cells. Panel A), upper plot, Phospho-(Ser) PKC substrate MFI in the indicated (white line) cell ROI was calculated as described in Materials and Methods for C3 control and P5 clones expressing HA-YFP-FMNL1β, either non-stimulated (Cont) or PMA-stimulated. The red line indicates the average Phospho- (Ser) PKC substrate MFI for each group. In the lower panel, left side, representative AIPs of the indicated channels of the transfected clones, stimulated with PMA, are shown. In the right side, quantification of Phospho-(Ser) PKC substrate MFI in C3 control and P5 clones expressing HA-YFP-FMNL1β, either non-stimulated (Cont) or PMA-stimulated, represented as fold induction of Phospho-(Ser) PKC substrate MFI (mean±SD), summarizing the results obtained in several (n=4) experiments. Panel B), upper diagram, calculation of the Phospho-(Ser) PKC substrate MFI in the specified (continuous white line) cell ROI was calculated as indicated in Materials and Methods, for C3 control and P5 PKCδ-interfered clones expressing HA-YFP-FMNL1β either non-stimulated (Cont) or IS-stimulated. The red line indicates the average Phospho- (Ser) PKC substrate MFI for each group. In the lower left panel, representative AIPs of the indicated channels of the IS-stimulated transfected clones (Raji cells are labelled with a white discontinuous line) are show. In the right side, quantification of Phospho- (Ser) PKC substrate MFI in C3 control and P5 PKCδ-interfered clones expressing HA-YFP-FMNL1β, either non-stimulated (Cont) or IS-stimulated, represented as fold induction of Phospho-(Ser) PKC substrate MFI (mean±SD), and summarizing the results obtained in several (n=5) experiments. NS, not significant; **, p≤0.05. Panel C), IS conjugates made by C3 control and P5 PKCδ-interfered clones expressing HA-YFP-FMNL1β were imaged by confocal microscopy to analyse the colocalization of HA-YFP-FMNL1β and Phospho-(Ser) PKC substrate signals, as indicated in Materials and Methods. In the upper row, MIPs of the indicated channels are shown. The white rectangles enclose the IS ROIs that were used for colocalization analyses in the focal planes indicated in the lower row. Colocalization pixels are shown in white colour on the colocalizations masks, as well as the corresponding Pearson’s coefficients. The lower plot shows the Pearson’s coefficients corresponding to several analyses similar to the one shown in the upper row, for synapses made by C3 and P5 clones expressing HA-YFP-FMNL1β or HA-YFP-FMNL1α. The red line indicates the average Pearson’s coefficient for each cell group. NS, not significant.

### PKCδ induces paxillin phosphorylation at the MTOC

The results from the previous section suggest that PKCδ-mediated FMNL1β phosphorylation at the IS may underlie F-actin reorganization at the IS, contributing to MTOC polarization. Although PKCδ appears to regulate centrosomal area F-actin (Fig. 2C), FMNL1β does not appear to participate in this regulation (Fig. 5C). A possible downstream effector of PKCδ in centrosomal area F-actin reorganization could be the actin regulatory protein paxillin, whose phosphorylation at threonine 538 (T538) by PKCδ leads to the depolymerization of the actin cytoskeleton and regulates integrin-mediated adhesion and migration of B lymphoid cells [43]. Moreover, the MTOC cannot polarize to the IS in paxillin-interfered T lymphocytes [25]. In addition, paxillin phosphorylation is required for the degranulation of CTL [26]. To analyze the subcellular location of paxillin with respect to MVB/MTOC polarization we first co-expressed GFP-actin, CFP-CD63 and Cherry-paxillin in C3 control and P5 PKCδ- interfered Jurkat clones, challenged them with SEE-pulsed Raji cells and analyzed by time-lapse microscopy the subcellular localization of paxillin relative to the reorganization of cortical actin and MVB polarization upon IS formation. We observed that Cherry-paxillin exhibited a mainly diffuse cytosolic distribution as well as decorated punctate structures nearby the MVB (Suppl. Video 6). In the C3 control clone, these structures polarized together with MVB towards the IS, immediately after an intense and prolonged actin reorganization at the IS [14] (Suppl. Video 6). In contrast, in the P5 PKCδ-interfered clone, after a weak cortical actin reorganization at the IS, Cherry-paxillin in punctate structures remained non-polarized together with CFP-CD63^+^ MVB (Suppl. Video 6). We next studied whether paxillin could be phosphorylated by PKCδ. When we challenged C3 control clone with SEE-pulsed Raji cells, immunofluorescence analysis showed that paxillin, colocalizing with MTOC (Fig. 7A and not shown) as previously described [44], was phosphorylated in T538 (Fig. 7A).

**Fig. 7.**
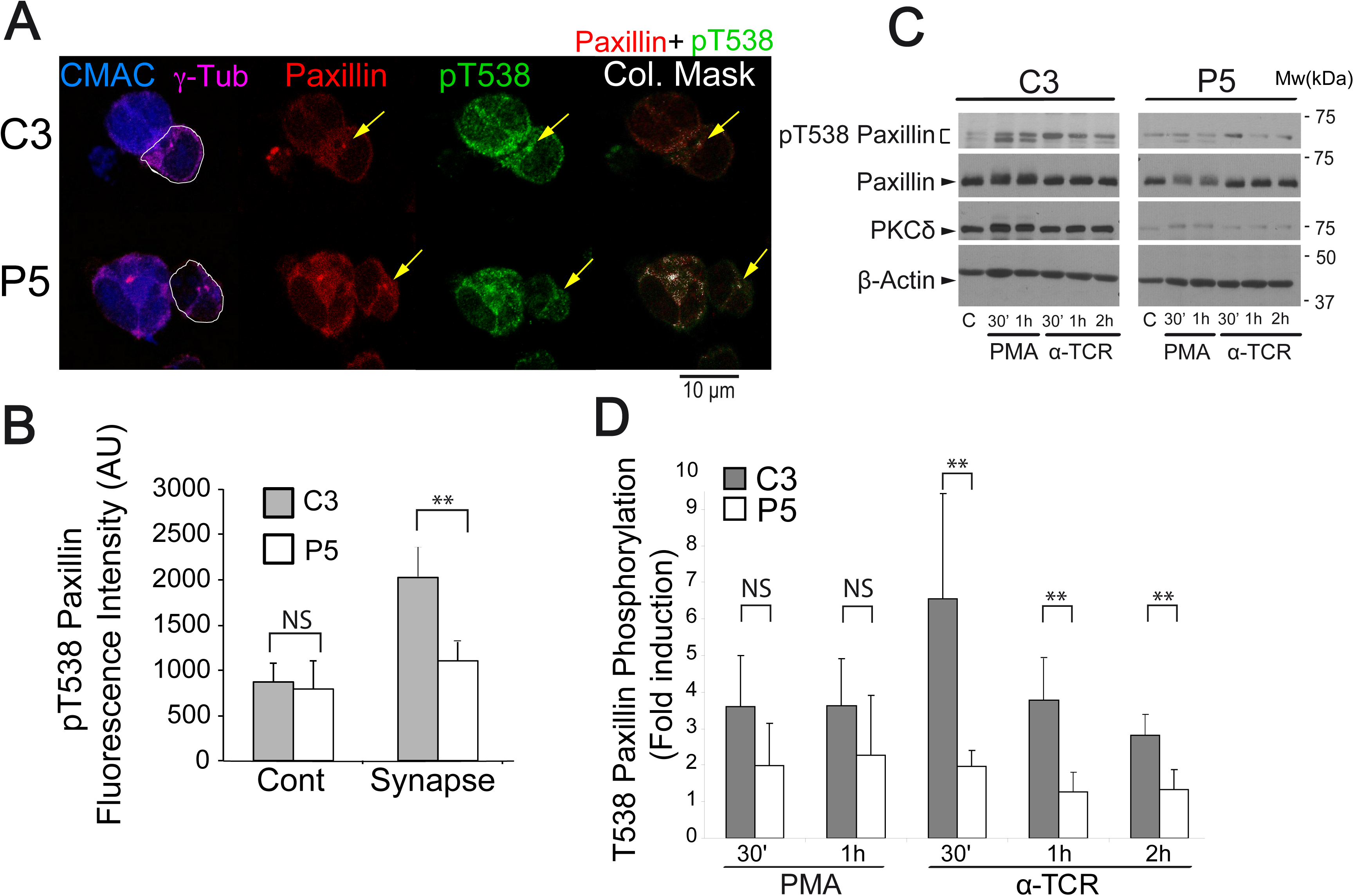
PKCδ regulates the phosphorylation of paxillin at T538. Panel A), C3 control and P5 PKCδ-interfered Jurkat clones were challenged with CMAC-labelled, SEE-pulsed Raji cells for 1 h, fixed, immunolabelled with anti-γ-tubulin, anti-paxillin and anti-phospho-T538 paxillin and imaged by confocal microscopy. White lines enclose Jurkat cells. In the right panels, colocalization masks of merged, paxillin and phospho-T538 paxillin channels, are represented in white. The yellow arrows indicate accumulations of paxillin and phospho-T538 paxillin. CMAC labelling of Raji cells in blue, γ-tubulin (MTOC) in magenta, paxillin in red and phospho-T538 paxillin in green. Panel B), C3 control and P5 PKCδ-interfered Jurkat clones were either untreated (Cont) or challenge with SEE-pulsed Raji cells (Synapse) and immunolabelled as in A. Phospho-T538 paxillin signals were internally normalized to paxillin signals as described in Materials and Methods. Panel C), C3 control and P5 PKCδ-interfered Jurkat clones untreated (C), or stimulated with either PMA or plastic-bound anti-TCR, were lysed at the indicated times. Cell lysates were analyzed by WB with antibodies against phospho-T538-paxillin, paxillin, PKCδ and β-actin. Panel D), Fold induction of paxillin T538 phosphorylation, normalized to paxillin levels, was evaluated by WB quantification of several experiments similar to that described in panel C. Data are means plus SD (*n* = 3). NS, not significant; **, p≤0.05.

To analyze phosphorylation of paxillin, we carried out image quantitation analysis and found that the pT538-paxillin MFI signal induced upon IS formation was significantly higher in the C3 control clone than in the P5 PKCδ-interfered clone (Fig. 7B). Since Raji cells express very high levels of paxillin (not shown), WB analysis of synaptic conjugates would not allow distinguishing Raji-derived paxillin from Jurkat clones-derived paxillin. Thus, to quantitate the relative, T538 paxillin phosphorylation by WB we stimulated control and PKCδ-interfered Jurkat clones with PMA or plastic-bound anti-TCR. An increase in pT538-paxillin signal was observed after stimulation of C3 control clone with PMA or anti-TCR (Fig. 7C, 7D), which was partially associated with a mobility shift of paxillin. However, the normalized induction of phosphorylation in T538 paxillin by anti-TCR stimulation in the P5 PKCδ-interfered clone was significantly lower than in the C3 control clone (Fig. 7D). Furthermore, GFP-PKCδ expression in P5 PKCδ-interfered clone stimulated with anti-TCR for 1 h raised T538 paxillin phosphorylation to the levels found in C3 control clone expressing GFP-PKCδ (Suppl. Fig. S11). Together these data indicate that PKCδ regulates phosphorylation and hence probably paxillin subcellular location and function during IS formation.

## DISCUSSION

We have identified here for the first time a positive regulatory role of PKCδ acting on both FMNL1β and paxillin-controlled actin regulatory pathways that govern MTOC/MVB polarization to the IS in Th lymphocytes. FMNL1β and paxillin, most probably, coordinately control F-actin reorganization at the IS and around the MTOC, respectively, and may collaborate in MTOC polarization and subsequent exosome secretory traffic [14]. PKCδ is known to regulate polarized secretion in T lymphocytes [14, 37, 45], and directional migration [46] [47], but no studies regarding the molecular bases controlling the PKCδ effect on polarized traffic have been reported to date. The spatial and temporal control of F-actin polymerization is fundamental for many cellular processes including cell migration, division, vesicle trafficking, and response to agonists. In this context, T and B lymphocytes are the only cells in which agonists, namely antigens, by binding to their TCR and BCR at the IS, induce polarized MVB traffic [48] and exosome secretion [13, 49] [15, 17]. This inducible polarized secretory traffic is controlled by actin cytoskeleton reorganization [5] [14]. In this paper we have dissected some mechanisms by which two distinct networks of F-actin (cortical F-actin at the IS and centrosomal area F-actin) are reorganized and their contribution to the processes of MTOC and MVB polarization upon TCR activation at the IS.

The understanding of the diverse molecular mechanisms controlling the distinct F-actin networks is an important and general biological issue [50, 51]. However, several contradictory results exist regarding the relative contribution of the different F-actin-regulatory pathways and the distinct F-actin networks to cell polarity in general [52] [53], and to MTOC polarization and subsequent secretion in T lymphocytes in particular [23] [7] [5]. Thus, while some results suggest (but do not formally demonstrate) that the cortical actin reorganization at the IS is necessary and sufficient for MTOC/lytic granules and cytokine-containing granules polarization [5] [7] [22], several other evidences show that the two major F-actin-regulatory pathways (Arp2/3 acting on cortical F-actin network and FMNL1 on non-cortical F-actin network) can independently act in the regulation of MTOC polarization [23]. In this context, it has been shown that MTOC is a F-actin organizing center [27] and centrosomal F-actin depletion appears to be crucial to allow MTOC polarization towards the IS in B lymphocytes stimulated with BCR-ligand-coated beads [28]. In the same B lymphocyte model, F-actin depletion around the MTOC, F-actin reorganization at the IS, and MTOC polarization depend on proteasome activity [29]. These data do not allow inferring either the sufficiency or the relative contribution of each F-actin network (cortical and centrosomal) to MTOC polarization. In addition, the results of this B lymphocyte model cannot be directly extrapolated to T lymphocytes forming cell-to-cell conjugates as used here.

The above-mentioned results on B lymphocytes, together with our previous observation regarding the effect of PKCδ on cortical F-actin on the IS, prompted us to study the involvement of centrosomal area F-actin in MTOC polarization in cell-to-cell synapses made by T lymphocytes and its potential regulation by PKCδ. We found a statistically significant negative correlation between MTOC polarization and centrosomal area F-actin (Fig. 2B, please compare with Fig. 5g from ref. [28]), and significant differences in centrosomal area F-actin among control and PKCδ-interfered clones (Fig. 2C). The differences observed between the values of centrosomal area F-actin of control (0.93, C3; 0.99, C9) vs PKCδ-interfered clones (1.18, P5; 1.37, P6) (Fig. 2C) were similar to those caused by proteasome inhibition in B lymphocytes (1.00 in control vs 1.25 in proteasome inhibitor-treated cells), that inhibited MTOC polarization [29]. Thus, it is conceivable that the centrosomal area F-actin changes we observed, alone or in combination with the alterations in cortical F-actin at the IS, may in part underlie the deficiency in MTOC polarization in PKCδ-interfered clones (Fig. 1B. The diameter determining the centrosome volume we have used (2 μm) to measure centrosomal area F-actin includes the wide range of centrosome radius from prophase to metaphase (400-2000 nm) [54]. Thus, this volume centered at the MTOC can include all the potential oscillations in centrosome radius, which indeed were noticeable in some of our results when a pericentriolar marker (DsRed2Centrin) was utilised (Fig. 2D, lower panels). Although some authors have employed the same ROI size to measure centrosomal F-actin changes in polarizing lymphocytes [28], it cannot be excluded that other organelles reorganizing F-actin or tubulin cytoskeleton may be included in this area, such as the Golgi [30] or endosomes/MVB [55] [56]. Among these organelles, we can rule out the contribution of F-actin at the Golgi to the observed MTOC polarization, since re-expression of FMNL1γ, which is the only FMNL1 isoform capable to regulate F-actin reorganization at the Golgi [30], was unable to rescue both centrosomal area F-actin and MTOC polarization to control levels (Suppl. Figs. S7 and S8). Regarding the presence of MVB in the centrosomal area, our results show that MVB are included in the 2 μm MTOC–centered area (Fig. 1, Suppl. Video 1 and Supp. Fig. S1), and thus we cannot exclude the contribution of MVB to F-actin reorganization at the centrosomal area during centrosome reorientation. Since MVB and MTOC center of masses are coincident and both organelles polarize (or not) together to the IS (Suppl. Fig. S1), the molecular mechanisms controlling these processes are, most probably, common for both simultaneously polarizing organelles. At this stage, we can only speculate about the necessity of the two F-actin reorganization processes or the sufficiency of one of these for MTOC polarization. In this context, FMNL1-interfered cells re-expressing FMNL1β exhibited high centrosomal area F-actin, polarized MTOC, and a significant F-actin depleted area at the central IS (Fig. 5, Suppl. Fig. S9). This suggests that, when only the FMNL1β isoform is present, centrosomal area F-actin disorganization is not necessary, and F-actin reorganization at the IS is sufficient, for MTOC polarization. In addition, the fact that neither FMNL1α nor FMNL1γ alone were capable to rescue both MTOC polarization (Suppl. Fig. S7) and centrosomal area F-actin (Suppl. Fig. S8) to control levels opens the possibility that any combination of FMNL1β with FMNL1α or FMNL1γ, or the three isoforms acting together, may collaborate on centrosomal area F-actin reorganization.

Regarding the contribution of the FMNL1 isoforms present in T lymphocytes (α, β and γ) [30] to the observed phenotype, and their potential role in PKCδ−dependent phosphorylation, it must be stressed that the anti-FMNL1 antibodies used for WB (C-5) and IP (clone A-4) recognized all these isoforms. In addition, the FMNL1-interfering vector transiently inhibiting the expression of all FMNL1 isoforms in Jurkat cells [30] decreased MTOC polarization to the levels observed in PKCδ- interfered clones (Fig. 5A, 5B). Thus, at this stage we cannot ascribe the observed effect of PKCδ on FMNL1 to the phosphorylation of one or several FMNL1 isoforms. Experiments with FMNL1 suppression, FMNL1 isoform-specific re-expression vectors [30] were performed to analyze the phosphorylation of the different FMNL1 isoforms and their specific role in MTOC polarization, as well as their subcellular localization. FMNL1β, but not FMNL1α or FMNL1γ, re-expression in YFP^+^ FMNL1-interfered cells restored MTOC polarization to that of control, YFP^-^ cells (Fig. 5A and Suppl. Fig. S7). In addition, we observed PMA- or IS formation-induced specific phosphorylation of FMNL1β, but not of FMNL1α, FMNL1γ or a C-terminal deletion mutant FMNL1 (FMNL1-⊗FH2)(Fig. 6B, Suppl. Fig. S10). The decrease in Phospho- (Ser) PKC substrate MFI signal induced by PMA stimulation occurring in HA-YFP-FMNL1α, HA-YFP-FMNL1γ and HA-YFP-FMNL1ΔFH2 (Suppl. Fig. S10B), but not in HA-YFP-FMNL1β-expressing cells (Suppl. Fig. S10A), when compared with non-transfected cells (Suppl. Fig. S10, compare panel A with panel B) is, most probably, caused by the interference in endogenous FMNL1β expression. FMNL1β possesses, but FMNL1α, FMNL1γ and FMNL1-⊗FH2 lack, a potential PKC phosphorylation site (S1086 in FMNL1β) in the DAD auto-inhibitory domain with high homology to that of FMNL2 (S1072) [42] [41] (Suppl. Fig. S6). The three FMNL1 isoforms share identical sequence from amino acid residue 1 to 1070, and diverge in the C-terminal region (Suppl. Fig. S6), which includes the DAD auto-inhibitory domain. PMA- and IS formation-induced FMNL1β phosphorylation, as well as IS formation-induced MTOC polarization, were decreased in P5 PKCδ-interfered clone (Fig. 6). Taken together, these results support that IS-induced, PKCδ−dependent phosphorylation in FMNL1β C-terminal region containing the auto-inhibitory domain (possibly at S1086) activates FMNL1β and mediates MTOC polarization. Further experiments re-expressing a mutated FMNL1β, non-phosphorylatable at S1086 (equivalent to FMNL2-S1072A described in [41]), are necessary to fully confirm this hypothesis and to study its role in MTOC polarization. In addition, it will be interesting to analyse whether a non-phosphorylatable FMNL1β mutant can rescue the relative F-actin-low cIS area as occurs with WT FMNL1β (Suppl. Fig. S9). FMNL1 or PKCδ interference inhibited MTOC polarization to similar extent (Fig. 5B), suggesting that PKCδ and FMNL1 participate in the same F-actin regulatory pathway controlling MTOC polarization. In addition, FMNL1 interference in a PKCδ-interfered clone did not further decrease MTOC polarization (Fig. 5B), which is compatible with PKCδ and FMNL1 participating in the same F-actin regulatory pathway. Additional experiments will be necessary to formally address this point.

DAG and its negative regulator DGKα play essential roles in TCR-induced MVB and late endosomes polarized traffic and lytic granule secretion [57] [58, 59] [15, 16], but the molecular mechanisms involved remained largely unknown. We have previously established that PKCδ is necessary for MTOC/MVB polarization [14]. In addition, we have shown that TCR activation at the IS induces PKCδ activation, which appears to be mediated by DAG-induced, PKCδ recruitment to MVB endomembranes [14]. We show here that upon IS formation activated PKCδ, directly or indirectly, induces FMNL1β phosphorylation at its C-terminal, autoregulatory region, possibly at S1086, although this has not been demonstrated yet. This may release FMNL1β from its C-terminal autoinhibitory domain, rendering FMNL1β capable of reorganizing F-actin at the IS. In addition, FMNL1 colocalizes with F-actin at the IS but not with centrosomal area F-actin (Fig. 3), suggesting that FMNL1 mainly governs synaptic F-actin. Our results showing that PKCδ interference does not affect FMNL1 location in F-actin reorganization areas (Fig. 3) capable to regulate MTOC polarization [23, 28, 60] [5] [27] is consistent with the idea that postranslational modifications of FMNL1 such as phosphorylation, but not changes in FMNL1 subcellular location, may underlie PKCδ- mediated control of FMNL1 function.

Regarding the potential mechanisms involved in centrosomal area F-actin disassembly and MTOC polarization, PKCδ and paxillin colocalize in polarized T lymphocytes at or nearby the MTOC [46] [61] [44] [25, 26] [14], the location where we observed T538-phosphorylated paxillin (Fig. 7A). Paxillin and FMNL1β may thus coordinately regulate F-actin reorganization around the MTOC and at the IS, respectively. This possibility is supported by the fact that PKCδ interference decreases paxillin phosphorylation and enhances F-actin around the MTOC that, most probably, subsequently inhibits MTOC/MVB polarization (Fig. 1 and Suppl. Videos 2, 4 and 5) and exosome secretion at the IS [14] and AICD (Suppl. Video 4). PKCδ phosphorylates paxillin at T538 in vitro and in vivo, leading to F-actin depolymerization during integrin-mediated adhesion of a B cell line [43]. Furthermore, paxillin is constitutively associated with the MTOC in T lymphocytes [44] [26] and, upon target cell binding, with the peripheral SMAC (pSMAC) at the IS in CTLs [25]. Moreover, interfering with paxillin impedes MTOC polarization [25]. In CTLs, paxillin phosphorylation regulates the MTOC polarization to the IS [25]. However, we were unable to detect any paxillin accumulation at the IS when we observed the burst of F-actin at the IS (Suppl. Video 6, Fig. 7). Thus, in our Th model, PKCδ-dependent paxillin phosphorylation may govern F-actin reorganizations at locations different from the IS, such as around the MTOC, that may also contribute to the diminished MTOC polarization observed in PKCδ-interfered clones. This hypothesis is supported by the decrease of paxillin phosphorylation in PKCδ-interfered clones (Fig. 7) and its reversal by GFP-PKCδ expression (Suppl. Fig. S11), which also causes the recovery of the spatial and temporal reorganization of F-actin at the IS and MVB polarization to the IS [14]. This PKCδ-dependent, paxillin-regulated mechanism for centrosomal area F-actin reorganization appears to co-exist in T lymphocytes with the PKCδ-dependent, FMNL1β-regulated mechanism for cortical F-actin reorganization. More research is necessary (i. e., experiments involving a phospho-deficient mutant of paxillin at 538 residue) to establish the relative contribution of this mechanism to the polarization processes.

With respect to the biological significance of these finely-tuned and coordinated actin cytoskeleton regulatory mechanisms involved in MTOC polarization, it is remarkable that centrosomal area F-actin reorganization is triggered by antigen-receptor stimulation, which is a common event occurring both in B [28] and T lymphocytes (this paper) forming IS. In BCR-stimulated B lymphocytes, apart of F-actin reorganization at the IS [48], centrosomal F-actin-reorganization controls polarized and local secretion of lysosomes at the synaptic cleft that is involved in antigen extraction from APC, a crucial event involved in antigen processing and the acquisition of B cell-effector functions [48] [29] [62]. In T lymphocytes, a comparable centrosomal area F-actin reorganization regulates MTOC and MVB polarization (this paper) that eventually leads to secretory granule secretion (including exosomes) [14]. Remarkably, T and B lymphocytes share the ability to form IS with APC, a crucial event involved in the immune response upon antigen challenge via TCR or BCR stimulation [48] [63], and this induces exosome secretion [17, 64]. The IS is a highly dynamic and plastic signalling platform induced by antigen receptors stimulation and triggers exquisite mechanisms leading to optimal polarized and focused secretion at the synaptic cleft, to avoid the stimulation (or death) of bystander cells [65] [9] [48] [1] [35], or the non-specific extraction of antigens from APC [29] [62]. The existence in T lymphocytes of at least two PKCδ-controlled, coordinated regulatory mechanisms acting on two distinct F-actin networks, both controlling MTOC polarization and secretion, may establish subtle regulatory checkpoints to finely control IS-triggered polarized secretion, as previously suggested [23]. Polarized secretion guarantees the antigen specificity of the final response in many cell lineages of the immune system, including innate NK cells [66, 67], CTL [5, 38], B lymphocytes [29] [62], primary CD4+ T cells [68] and Jurkat cells [38]. The association of the absence of PKCδ with the generation of B and T lymphoproliferative disorders both in human and mice [69] [70], and the participation of PKCδ in the homeostasis of blood progenitors [71], support the important role of PKCδ in maintaining cell homeostasis. Further approaches are needed to extend our results regarding the contribution of PKCδ- dependent centrosomal area F-actin to polarized and focused secretion in directional and invasive cell migration, both of lymphoid and non-lymphoid cells. These approaches would clarify whether centrosomal area F-actin reorganization occurs only during IS formation or also participates in other biologically relevant cellular polarization processes.

## Acknowledgements

We are indebted and acknowledge Dr. D.D. Billadeau (Mayo Clinic, USA) for generous sharing of shFMNL1 and FMNL1 isoform rescue constructions. We acknowledge the excellent technical support from A. Sánchez and A. Garrido. We acknowledge Dr. A. Anel (Universidad de Zaragoza, Spain) for suggestions and critical reading of this manuscript and Dr. M.A. Alonso (CBM, CSIC) for reagents and scientific advice. Thanks to D. Morales (SIDI-UAM) and S. Gutiérrez (CNB, CSIC) for their superb expertise with confocal microscopy. This work was supported by grants from the Spanish Ministerio de Economía y Competitividad (MINECO), Plan Nacional de Investigación Científica (SAF2016-77561-R to M.I., which was in part granted with FEDER-EC-funding).

## Author Contributions Statement

V.C. and M.I. conceived and designed all the experiments. A.B., R.I., M.V., S.M., G.H., S.H. and V.C. did most of the experiments, analyzed data, and contributed to the writing of the manuscript. A.B., R.I. contributed to the MTOC/MVB polarization experiments and IS image analyses. A.B. performed the FMNL1 immunoprecipitation experiments. L.M. M.V. and S. M. contributed to the FMNL1 phosphorylation studies and image analyses, and M.V., S.M., J.B. and G.H. contributed to time-lapse studies and image analyses of area F-actin. A.S. performed the WB analysis and also contributed to paxillin phosphorylation experiments. M.I. conceptualized and coordinated the research, directed the study, analyzed data, and wrote the manuscript. All the authors contributed to the planning and designing of the experiments and to helpful discussions.

## Conflict of Interest

The authors report no conflict of interest.

Supplementary information is available at XXXX website.

## Supplementary Material

**Supplementary Fig. S1.**
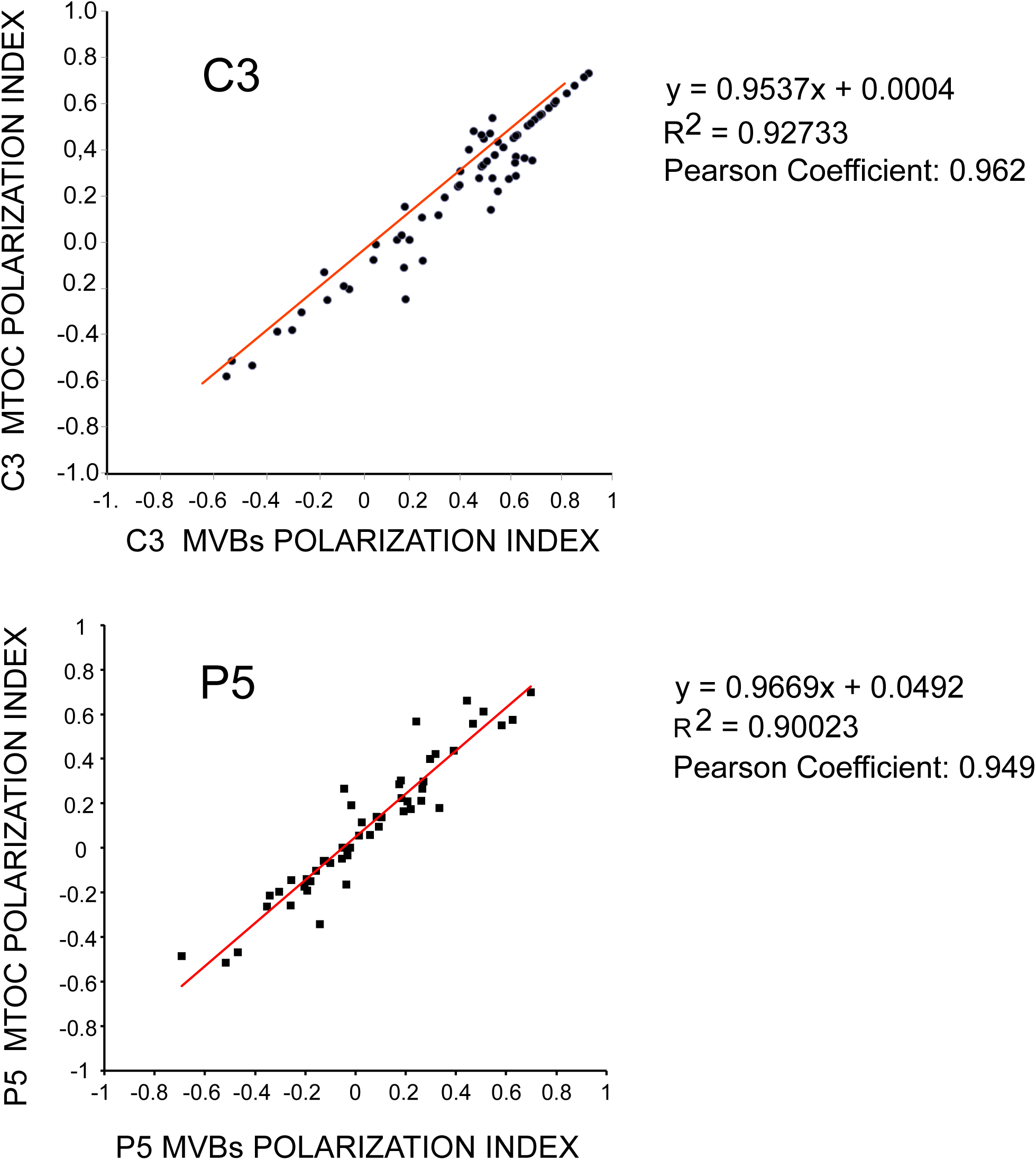
Correlation between MTOC and MVB polarization indexes. C3 control and P5 PKCδ-interfered clones were challenged with CMAC-labelled SEE-pulsed Raji cells for 1 h, fixed, stained with anti-γ-tubulin AF546 to label MTOC and anti-CD63 AF647, to label MVB and imaged by confocal fluorescence microscopy. Subsequently, MTOC and MVB Polarization Indexes were calculated for each cell, for both clones forming synapses, as described in Materials and Methods and Fig. 1A, and represented. Linear correlation analyses between the two parameters for both clones are represented, as well as the corresponding linear correlation analysis data and the Pearson’s coefficients. Red line represents the adjusted, regression line. Results are representative of the data from several experiments (n=3) with similar results. This figure is related to Fig. 1A.

**Supplementary Fig. S2.**
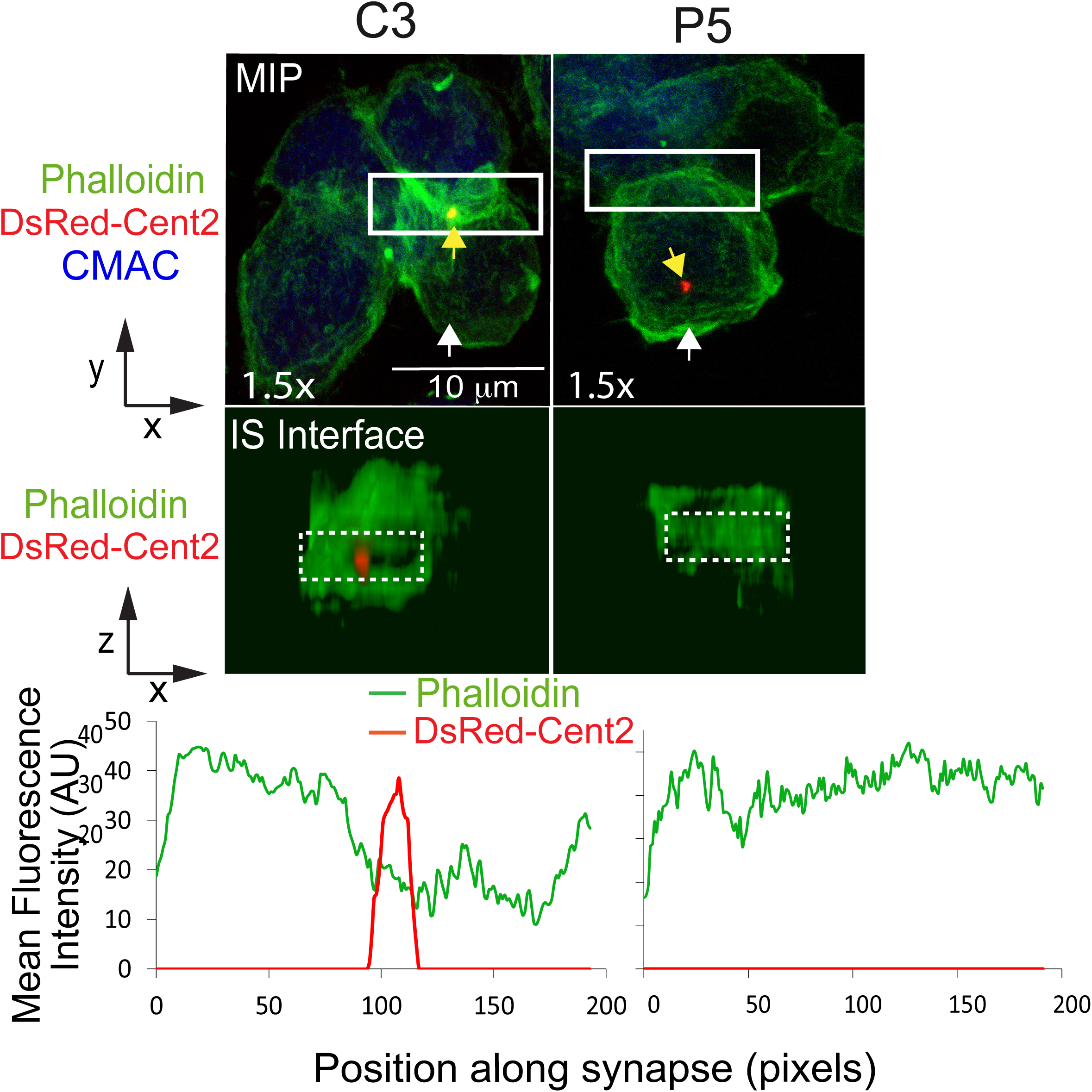
PKCδ regulates the polarization of the MTOC to the F-actin depleted area at the cIS. C3 control and P5 PKCδ-interfered clones expressing dsRed-Cent2 were challenged with CMAC-labelled SEE-pulsed Raji cells for 1 h, fixed, stained with phalloidin AF488 and imaged by confocal fluorescence microscopy. Upper panels: top views correspond to the Maximal Intensity Projection (MIP) of the indicated, three merged channels, in a representative example. White arrows indicate the direction to visualize the face on views of the synapse (IS interface) enclosed by the rectangular ROIs (white line), whereas yellow arrows indicate the MTOC position. Lower panels: face on views of the IS. The enlarged ROIs from the upper panel (1.5x zoom) were used to generate the IS interface images shown in the lower panels, as shown in Fig. 1C and Suppl. Video 3. The IS interface images of both the phalloidin and dsRed-Cent2 merged channels are represented. Lower diagrams: phalloidin and dsRed-Cent2 MFI vs position along the indicated, rectangular ROIs (white discontinuous line) embed in the face on views from the lower panels. CMAC labelling of Raji cells in blue, phalloidin in green and dsRed-Cent2 in red. Scale bars, 10 µm. Data are representative of the results obtained in 35 synapses. This figure is related to Fig. 1C and Suppl. Video 3.

**Supplementary Fig. S3.**
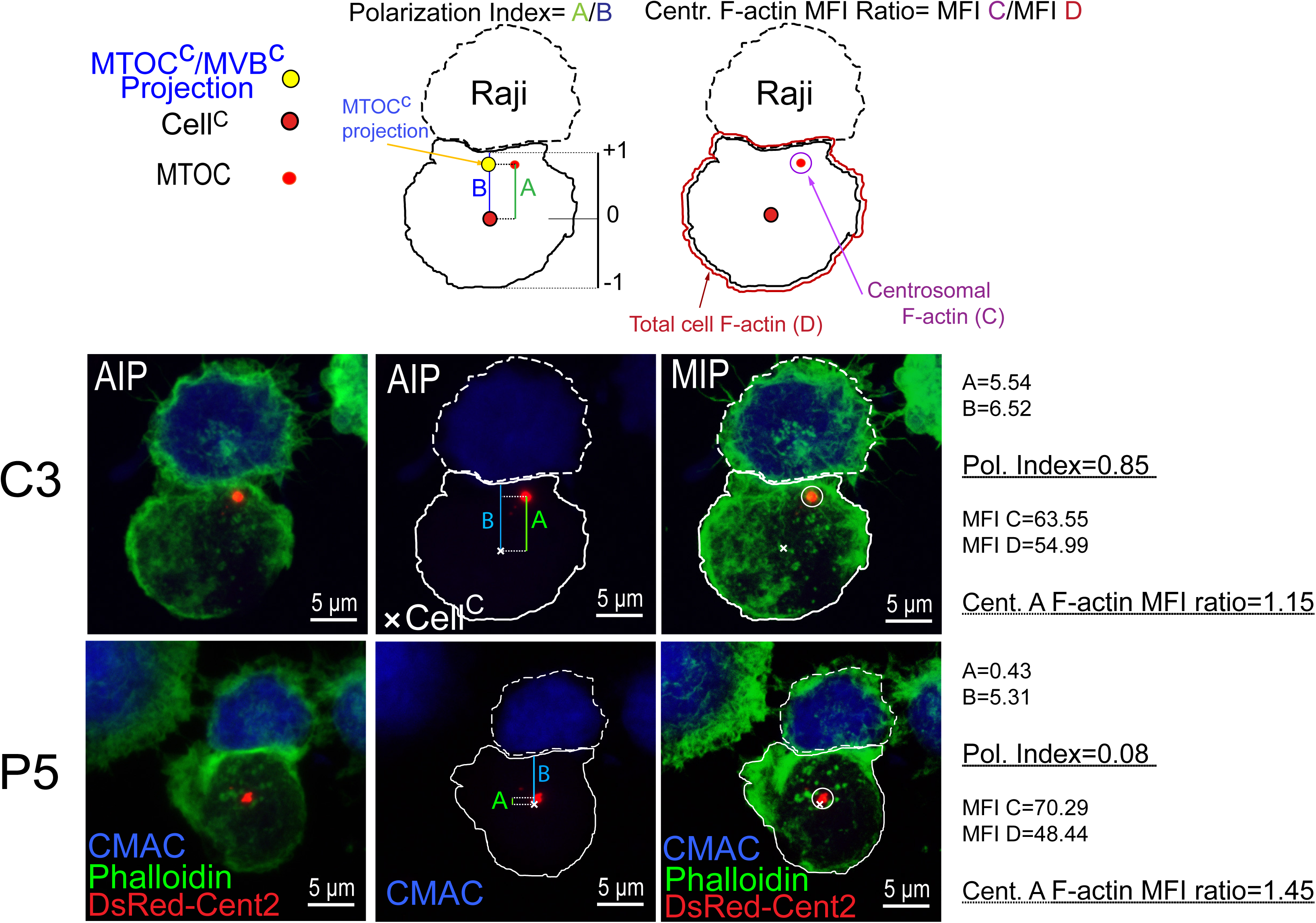
PKCδ regulates both the polarization of the MTOC and the centrosomal area F-actin. C3 control and P5 PKCδ-interfered clones expressing dsRed-Cent2 were challenged with CMAC-labelled SEE-pulsed Raji cells for 1 h, fixed, stained phalloidin AF488 to label F-actin, and imaged by confocal fluorescence microscopy. In the upper scheme, the distances in color (A, green; B, blue) used for the calculation of Pol. Index (A/B) are indicated in the diagrams, as in Fig.1A. In addition, the 2 μm diameter centrosomal area ROI (small white line circle) and the total cell ROI (continuous white line) that were used to calculate the centrosomal area F-actin MFI ratio (MFI C/MFI D), as shown in Fig. 2A, are represented. In the lower panels, AIP (left and center) and MIP (right) of the indicated, merged channels of representative images for both C3 and P5 forming synapses are shown, including the superimposed diagrams in the central panel. Superimposed, white crosses label the Cell^c^. The Raji cells and the Jurkat clones are labelled with discontinuous and continuous white lines, respectively. On the right side, the polarization index and the centrosomal area F-actin MFI ratio corresponding to these synapses are shown. Data are representative of the results obtained in 28 synapses. This figure is related to Figs. 1 and 2.

**Suppl. Fig S4.**
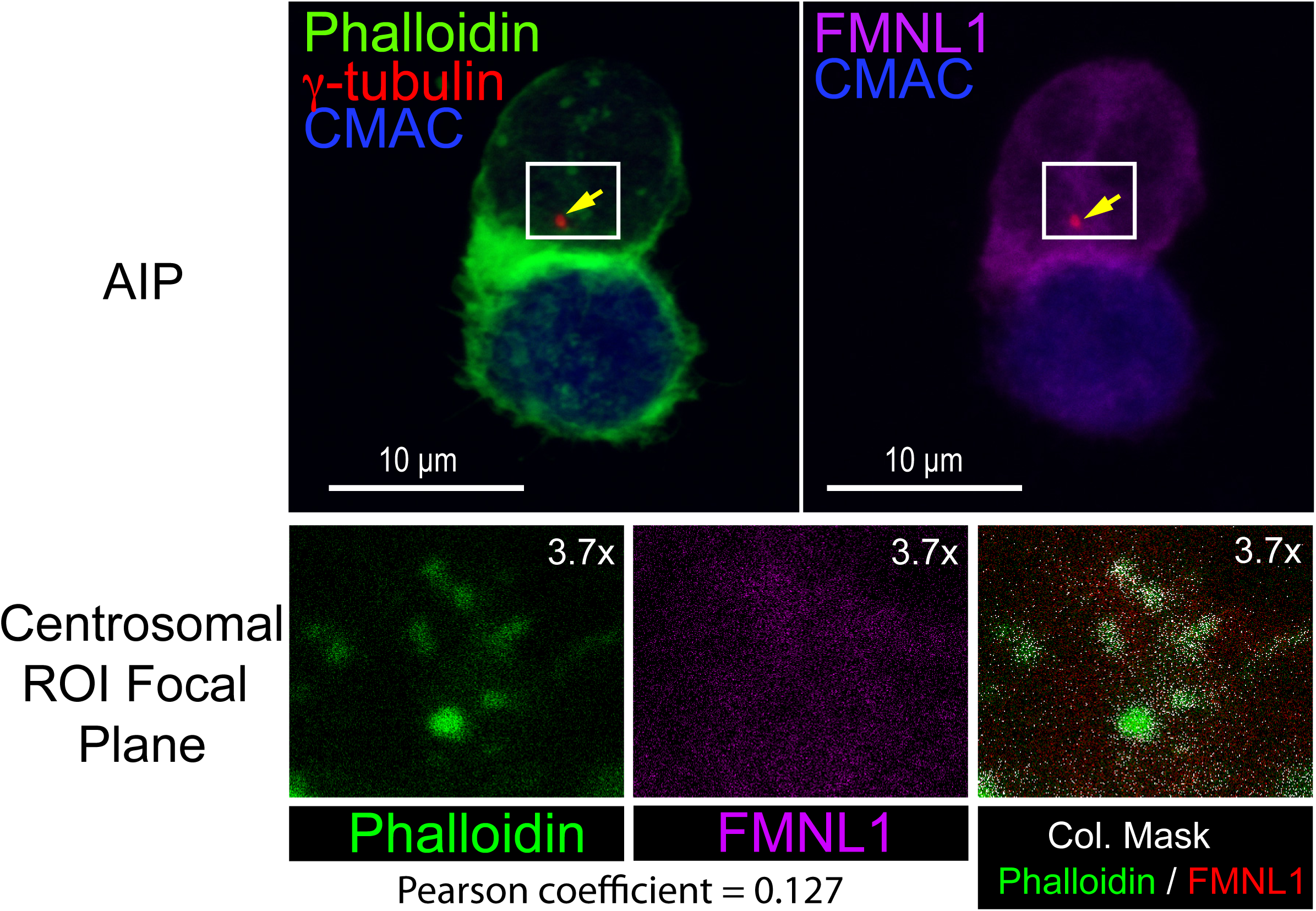
Subcellular localization of FMNL1 and MTOC. C3 control clone was challenged with CMAC-labelled SEE-pulsed Raji cells for 1 h, fixed, stained with anti-FMNL1 AF546 and phalloidin AF488 to label F-actin, and imaged by confocal fluorescence microscopy. In the upper panels, representative AIPs of the indicated, merged channels of synaptic conjugates made by C3 control clone are shown. White rectangles enclose the centrosomal (MTOC) ROIs used for the colocalization analyses of the indicated channels in optical sections, shown in the lower row of panels. On the right side, colocalization mask corresponding to merged, phalloidin and FMNL1 channels, is shown in white. Pearson’s coefficient corresponding to the colocalization analysis is indicated. Yellow arrow labels the MTOC. This figure is related to Fig. 3 in the manuscript.

**Suppl. Fig S5.**
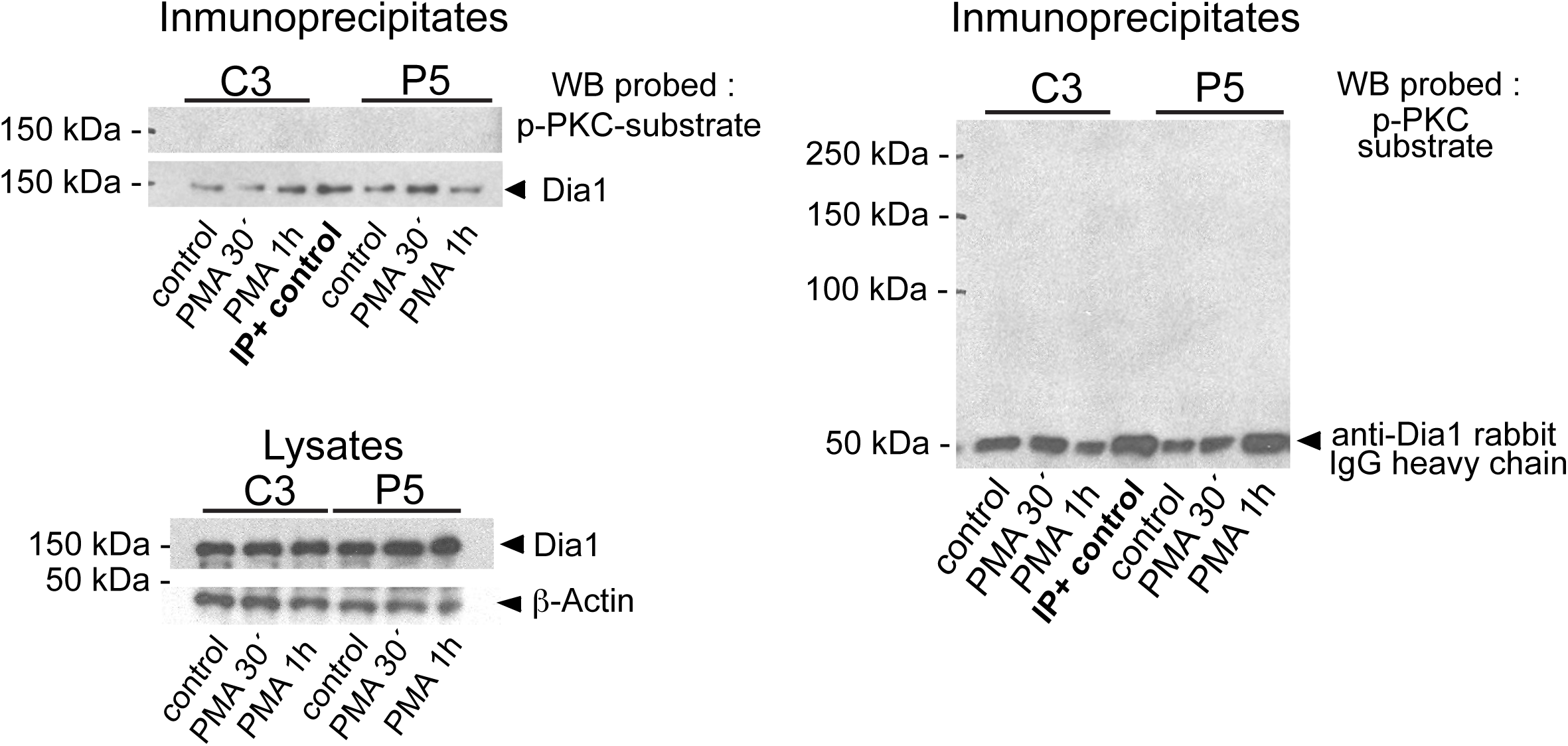
Dia1 is not phosphorylated upon PKC stimulation. C3 control clone was untreated or stimulated with PMA and, subsequently, cells were lysed at the indicated times. In the left panels, the WB of anti-Dia1 immunoprecipitates (upper panel) and cell lysates (lower panel) were sequentially probed with ant-Phospho-(Ser) PKC substrate and anti-Dia1, to analyze the Phospho-(Ser) PKC signal and Dia1 content in the immunoprecipitates. In addition, in the right panel the immunoprecipitates were probed with Phospho-(Ser) PKC and then secondary, anti-rabbit-HRP to check the presence of rabbit polyclonal anti-Dia1 (immunoprecipitating antibody) in the immunoprecipitates and the secondary antibody. This figure is related to Fig. 4 in the manuscript.

**Suppl. Fig. S6.**
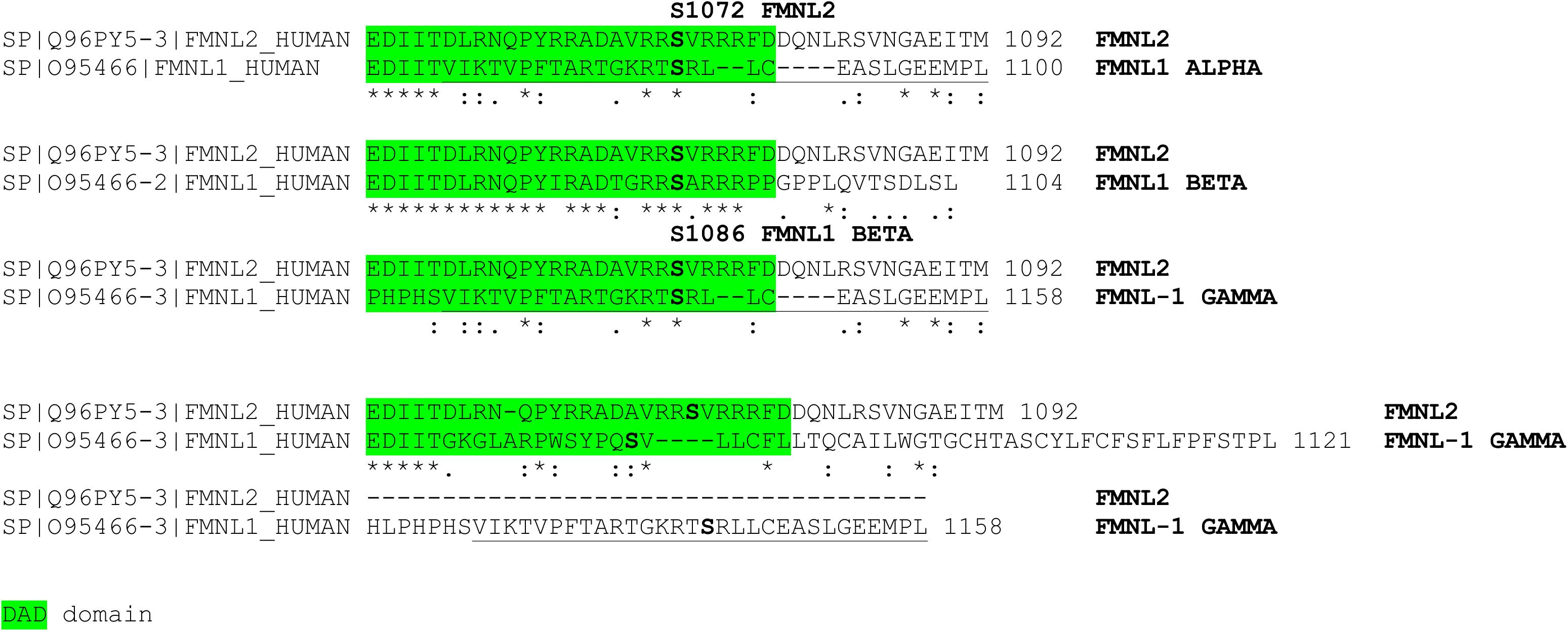
C-terminal alignment of FMNL1 isoforms. Amino acid sequences of the C termini of FMNL2 and FMNL1α and β, as well as of the isoform FMNL1γ containing a C-terminal intron retention, and sharing the final C-terminal amino acids with FMNL1α. The three FMNL1 isoforms share identical sequence from amino acid residue 1 to 1070, and diverge in the C-terminal region, which includes the DAD auto-inhibitory domain. The DAD domain sequence responsible for autoinhibition in the murine homolog is framed in green color, and the identical C-terminal amino acids in FMNL1α and FMNL1γ are underlined. A double alignment between FMNL2 and FMNL1γ is represented.

**Suppl. Fig. S7.**
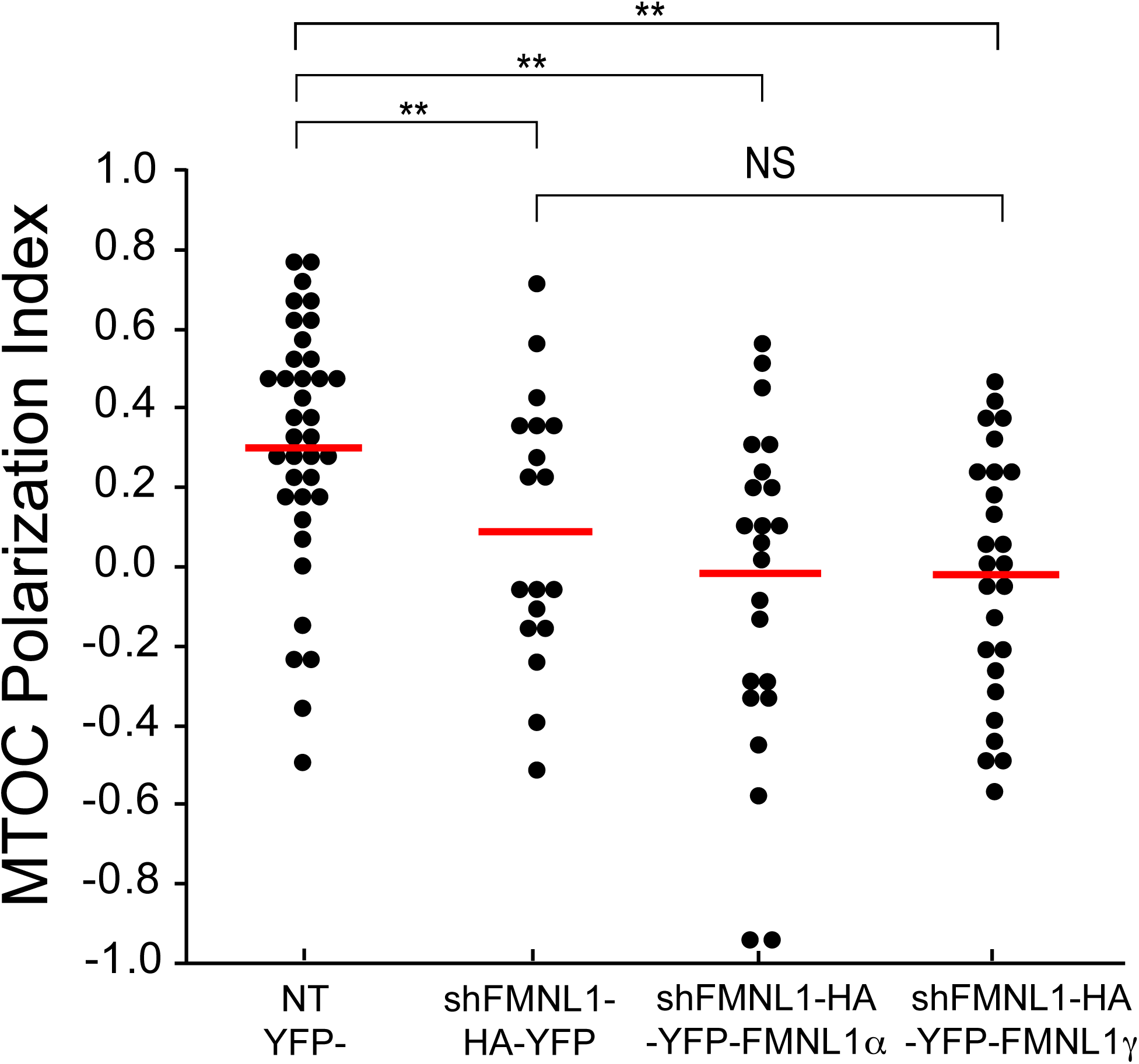
FMNL1α and FMNL1γ do not rescue MTOC polarization. C3 control was transfected with either control (shControl-HA-YFP), FMNL1 interfering (shFMNL1-HA-YFP), or FMNL1-interfering, YFP-FMNL1α expressing vector (shFMNL1-HA-YFP-FMNL1α) or FMNL1-interfering, YFP-FMNL1γ expressing vector (shFMNL1-HA-YFP-FMNL1γ). Subsequently, the transfected clones were challenged with CMAC-labelled SEE-pulsed Raji cells for 1 h, fixed, stained with anti-γ-tubulin AF546 (red) and imaged by confocal fluorescence microscopy. MTOC Pol. Index was calculated as indicated in Materials and Methods, for the indicated number of synaptic conjugates made by C3 control clone, transfected or not (NT YFP^-^ cells). This group includes all non-transfected cells from shFMNL1-HA-YFP, shFMNL1-HA-YFP-FMNL1α and shFMNL1-HA-YFP-FMNL1γ transfections as internal controls. Dot plot distribution and average Pol. Index (red horizontal line) are represented. NS, not significant. **, p ≤0.05. This figure is related to Fig. 5.

**Suppl. Fig. S8.**
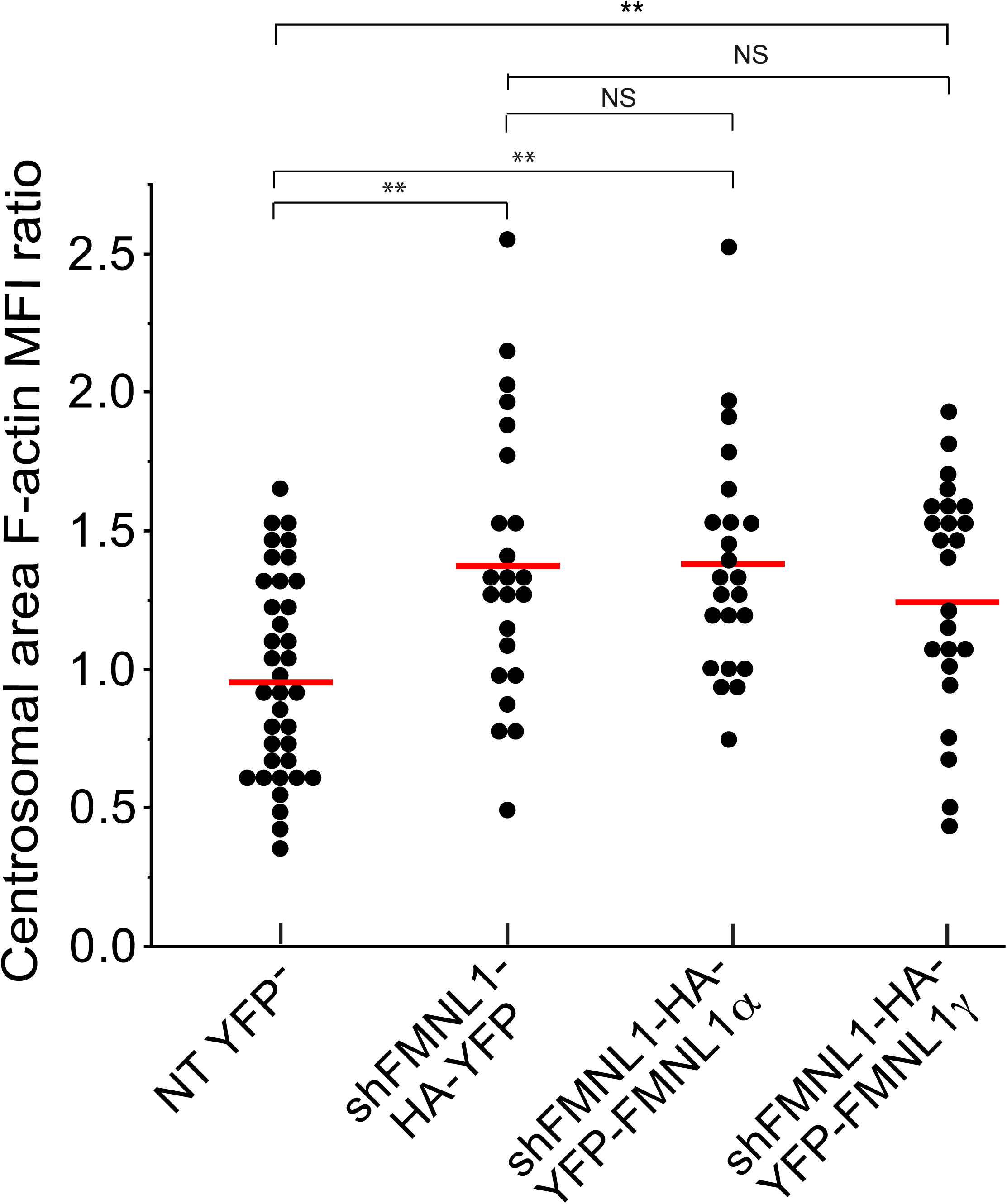
FMNL1α and FMNL1γ do not rescue centrosomal area F-actin reorganization. C3 control clone was transfected with either control (shControl-HA-YFP), FMNL1 interfering (shFMNL1-HA-YFP), or FMNL1-interfering, YFP-FMNL1α expressing vector (shFMNL1-HA-YFP-FMNL1α) or FMNL1-interfering, YFP-FMNL1γ expressing vector (shFMNL1-HA-YFP-FMNL1γ). Subsequently, the transfected clones were challenged with CMAC-labelled SEE-pulsed Raji cells for 1 h, fixed, stained with anti-γ-tubulin AF546 and phalloidin AF647 to label F-actin, and imaged by confocal fluorescence microscopy. Centrosomal area F-actin MFI ratio was calculated as indicated in Fig. 2 for the indicated number of synaptic conjugates made by C3 control clone, transfected or not (NT YFP^-^). The mean centrosomal area F-actin MFI ratio (red horizontal line) for each condition is represented. NS, not significant. **, p ≤0.05. This figure is related to Fig. 5C.

**Suppl. Fig. S9.**
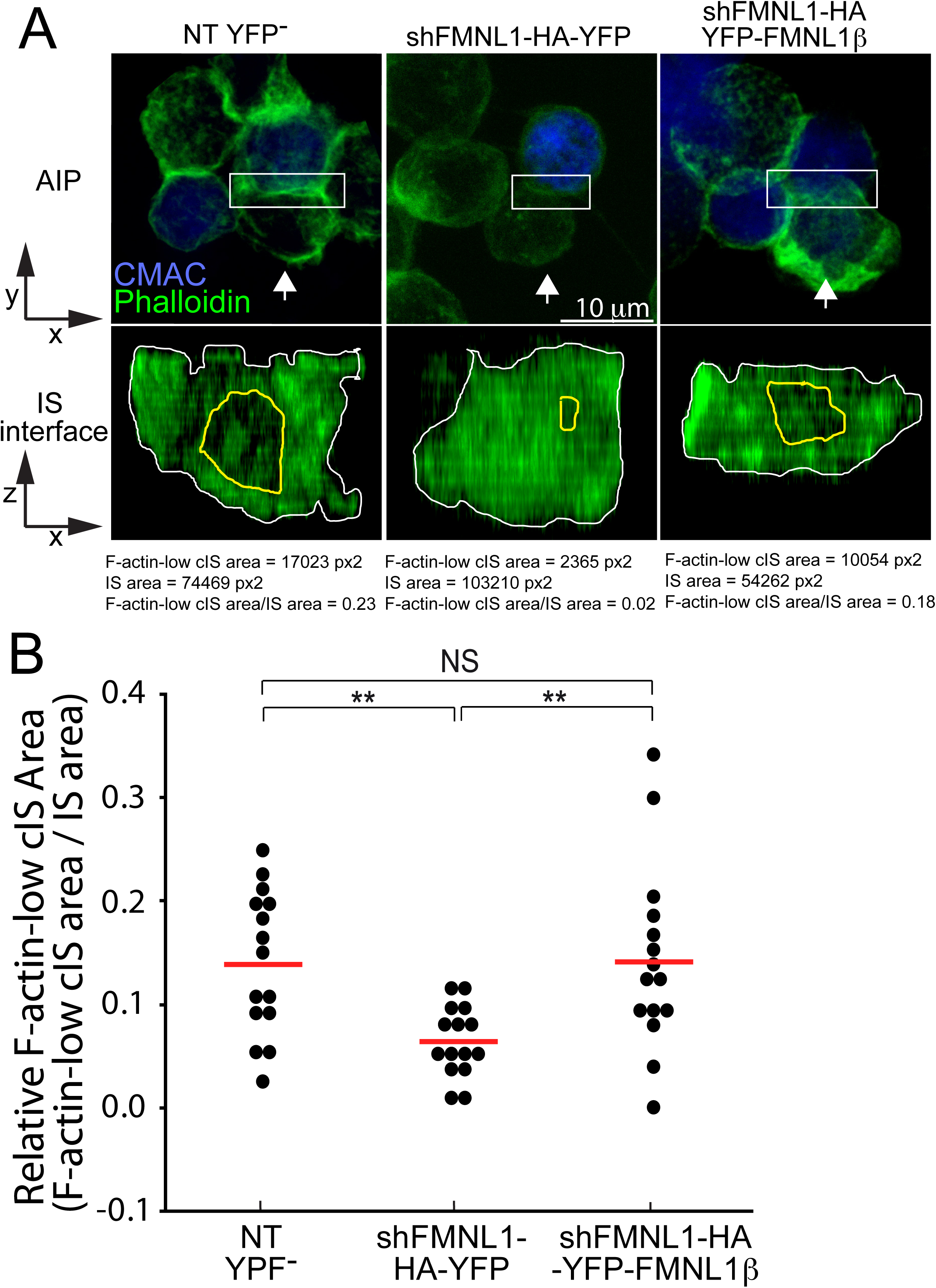
FMNL1β controls the reorganization of cortical F-actin at the immune synapse. C3 control clone was transfected with either FMNL1-interfering (shFMNL1-HA-YFP) or FMNL1-interfering, HA-YFP-FMNL1β expressing vector (shFMNL1-HA-YFP-FMNL1β). Subsequently, the cells were challenged with CMAC-labelled SEE-pulsed Raji cells for 1 h, fixed, stained with phalloidin AF488 (green) to label F-actin, and imaged by confocal fluorescence microscopy. Panel A), the upper panels include the top views correspond to the Average Intensity Projection (AIP) of the indicated, two merged channels of representative examples of non-transfected or transfected cells. White arrows indicate the direction to visualize the face on views of the synapse (IS interface) enclosed by the ROIs (white rectangles) as shown in Suppl. Video 3. In the lower panels, the enlarged ROIs (2x zoom) used to generate the IS interface images are shown. The areas of the F-actin-low region at cIS (Fact-low cIS area) (yellow line) and the synapse (IS area) (white line) were defined and measured as indicated in Materials and Methods, and the relative area of the F-actin-low region at the cIS (Fact-low cIS area / IS area) was calculated and represented. Panel B), relative area dot plot distributions and average area ratio (red horizontal lines) for the indicated number of IS conjugates made by non-transfected or transfected cells from one representative experiment out of 3 are showed. This figure is related to Suppl. Video 3. NS, not significant; **, p≤0.05.

**Suppl. Fig. S10.**
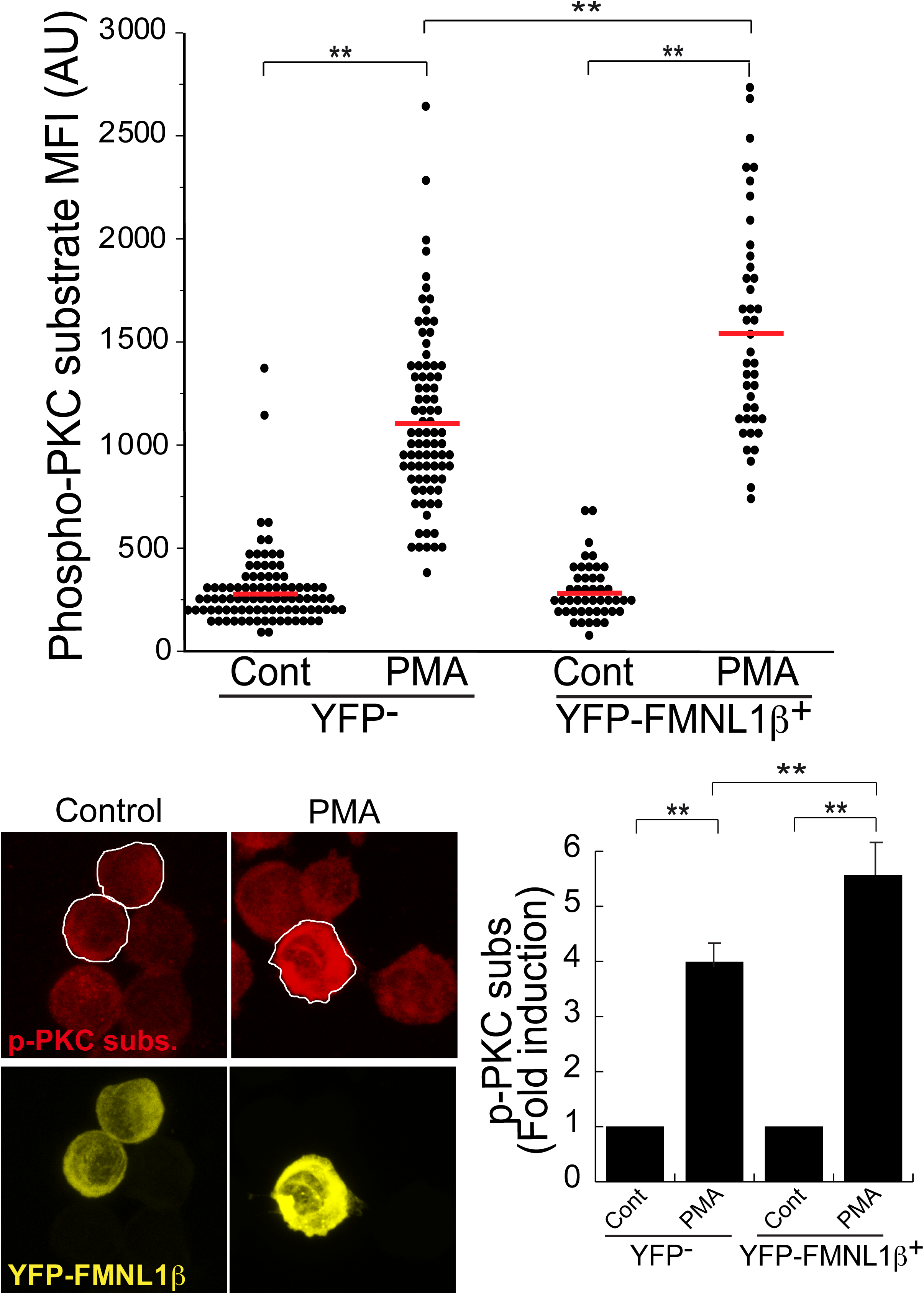
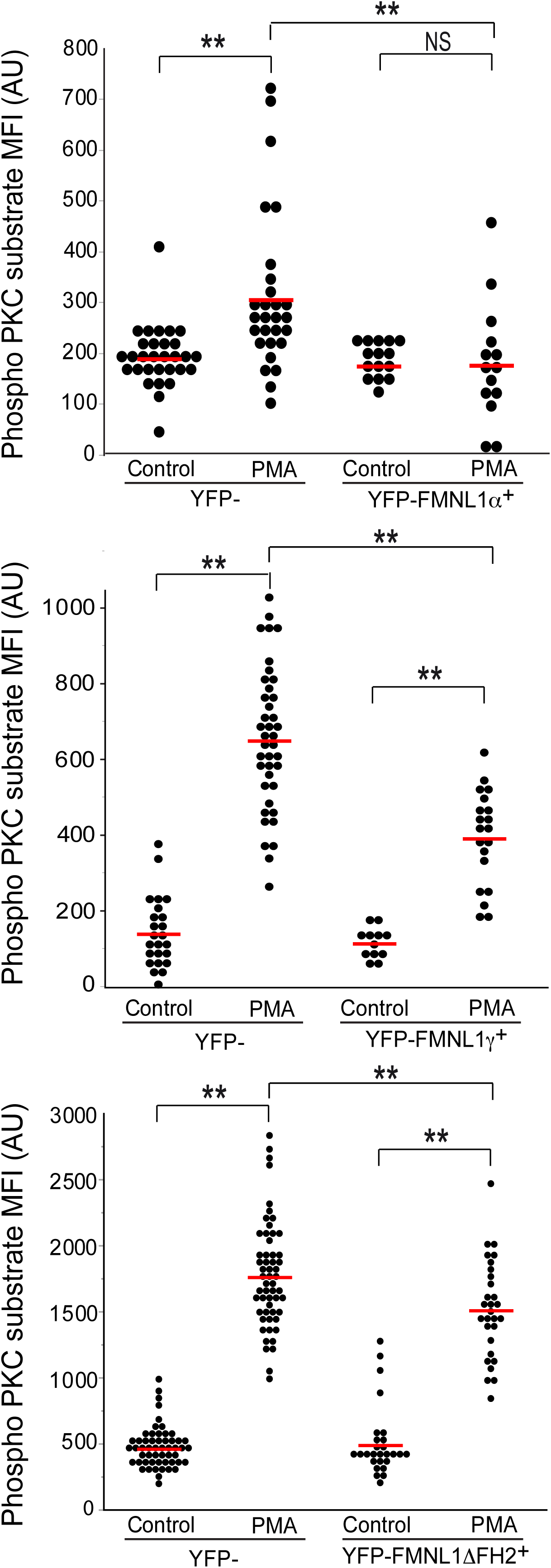
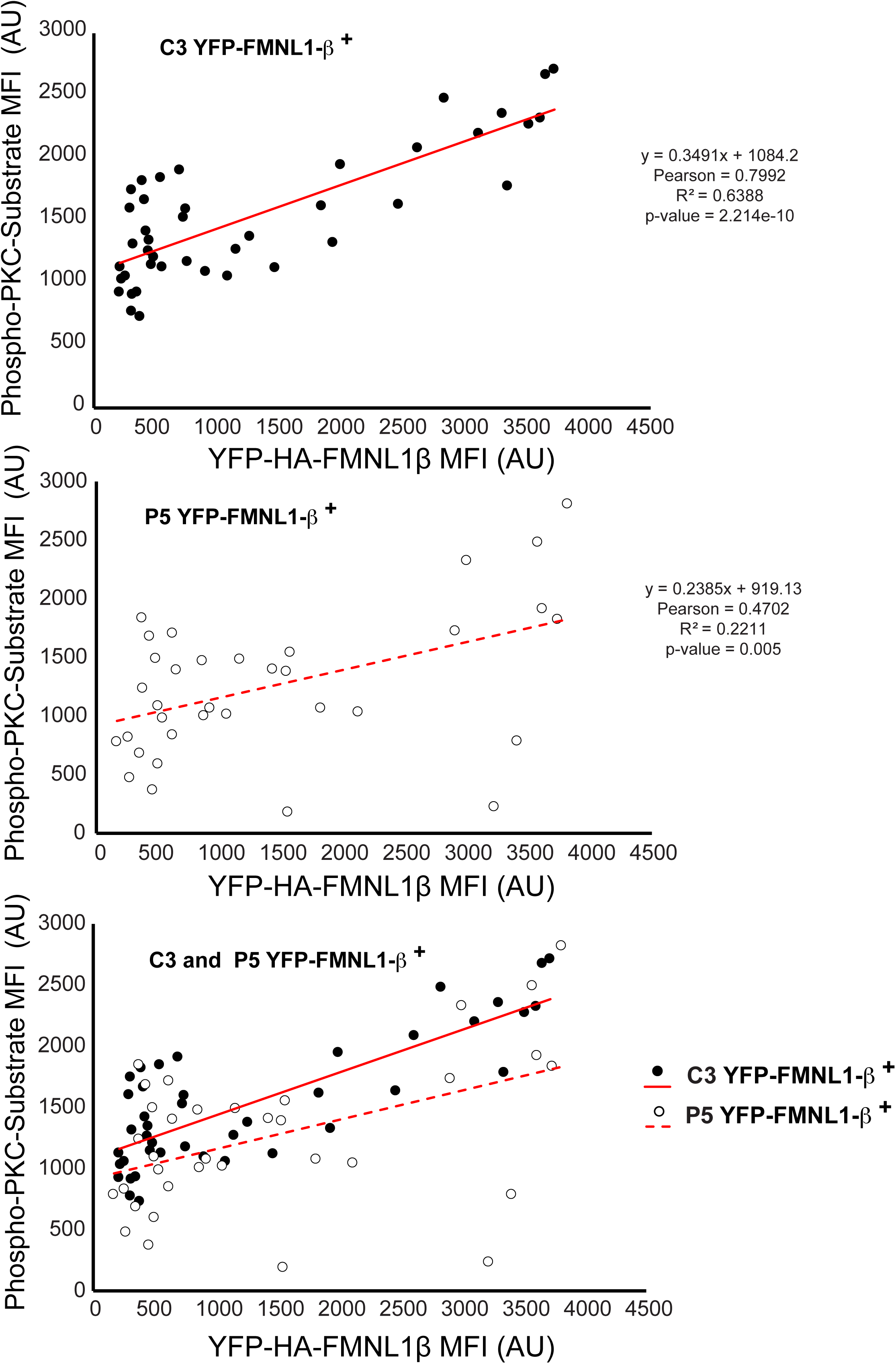
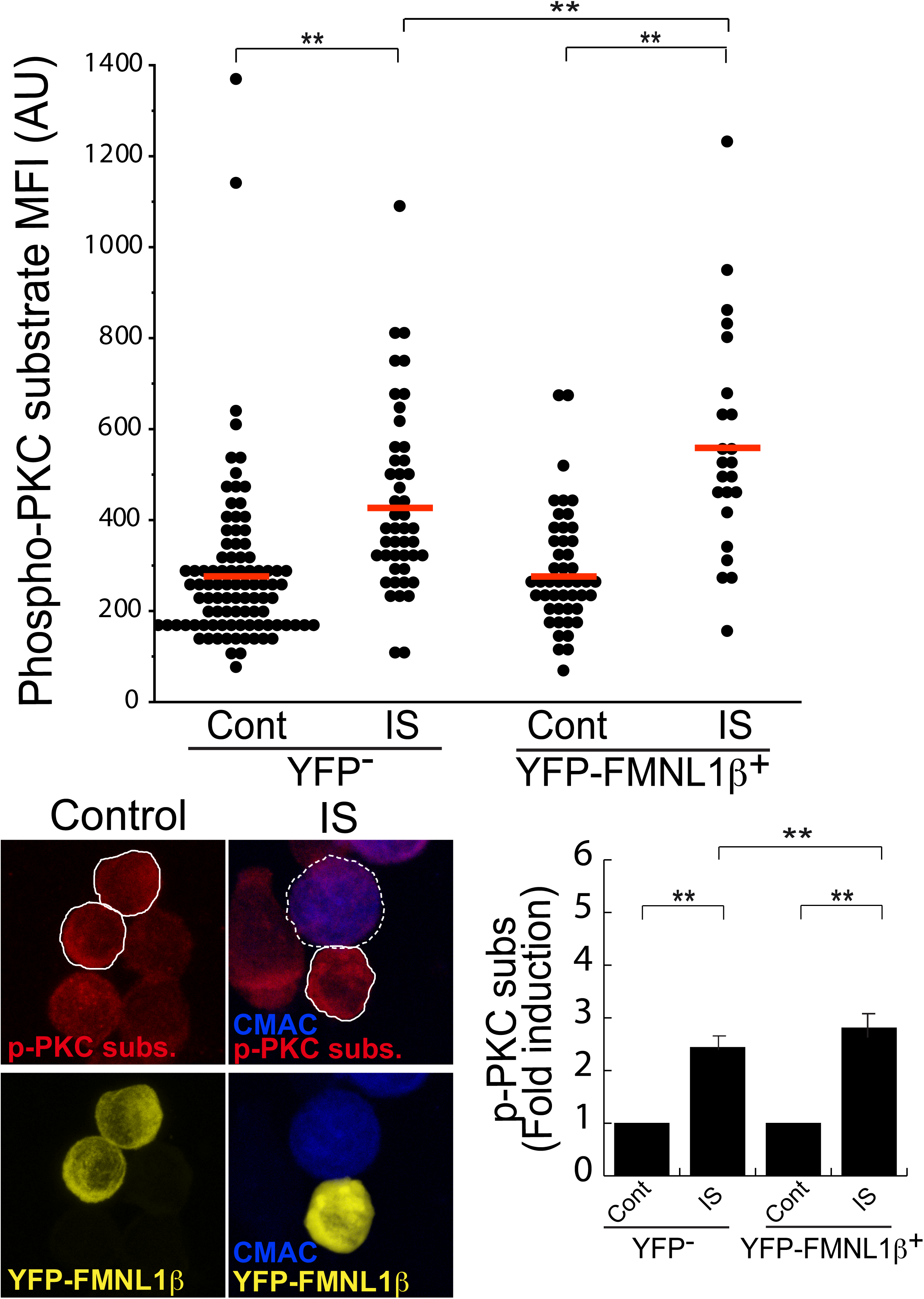
FMNL1β isoform is phosphorylated upon PKC stimulation with PMA. Panel A), C3 control clone was transfected with the FMNL1-interfering, HA-YFP-FMNL1β expressing vector (shFMNL1-HA-YFP-FMNL1β). Subsequently, the transfected clone was either untreated (Cont) or stimulated with PMA (30 min) fixed, stained with anti-Phospho-(Ser) PKC substrate AF647 (magenta) and imaged by confocal fluorescence microscopy. The upper plot shows Phospho-(Ser) PKC substrate MFIs of non-transfected (YFP^-^) or HA-YFP-FMNL1β-expressing (YFP-FMNL1β**^+^**), either non-stimulated (Cont) or PMA-stimulated (PMA), C3 cells, calculated using the cell ROI (white line in lower panels) as indicated in Materials and Methods. The red line indicates the average Phospho-(Ser) PKC substrate MFI for each group of cells. In the lower left panel, representative AIP confocal images of the indicated channels of the transfected cells, stimulated or not with PMA, are show. In the lower right plot, fold induction of Phospho-(Ser) PKC substrate MFI (mean+SD) in non-expressing (YFP^-^) or expressing HA-YFP-FMNL1β, either non-stimulated (Cont) or PMA-stimulated (PMA) cells, summarizing the results obtained in several (n=4) experiments. Panel B), same as panel A, but C3 control clone was transfected with the FMNL1-interfering, HA-YFP-FMNL1α, HA-YFP-FMNL1γ or HA-YFP-FMNL1ΔFH2 expressing vectors (shFMNL1-HA-YFP-FMNL1α, HA-YFP-FMNL1γ, shFMNL1-HA-YFP-FMNL1ΔFH2, respectively). The decrease in Phospho-(Ser) PKC substrate MFI signal induced by PMA stimulation occurring in HA-YFP-FMNL1α, HA-YFP-FMNL1γ and HA-YFP-FMNL1ΔFH2, but not in HA-YFP-FMNL1β-expressing cells (panel A), when compared with non-transfected cells is, most probably, caused by the interference in endogenous FMNL1β expression. Panel C), C3 control and P5 PKCδ-interfered clones expressing HA-YFP-FMNL1β were stimulated as in panel A, and imaged by confocal fluorescence microscopy to measure Phospho-(Ser) PKC substrate MFI and HA-YFP-FMNL1β MFI. Linear correlation analysis between these values corresponding to each individual cell is represented for each clone, and the Pearson’s correlation coefficients represented. Non-parametric Spearman’s correlation coefficients between these variables were ρ=0.6486 (p=1.354e-06) for C3, and ρ=0.4163 (p=0.015) for P5. Panel D), C3 control clone was transfected with the FMNL1-interfering, HA-YFP-FMNL1β expressing vector (shFMNL1-HA-YFP-FMNL1β). Subsequently, the transfected clone was either untreated (Cont), or challenged with CMAC-labelled SEE-pulsed Raji cells for 1 h (IS), fixed, stained with anti-Phospho-(Ser) PKC substrate AF647 (magenta) and imaged by confocal fluorescence microscopy. Yellow channel fluorescence identifies the HA-YFP-FMNL1β−expressing cells. The upper plot shows Phospho-(Ser) PKC substrate MFIs of non-transfected (YFP^-^) or HA-YFP-FMNL1β-expressing (YFP-FMNL1β**^+^**), either non-stimulated (Cont) or IS-stimulated (IS) cells, calculated using the cell ROI (white line in lower panels) as indicated in Materials and Methods. The red line indicates the average Phospho-(Ser) PKC substrate MFI for each group of cells. In the lower left panel, representative AIP confocal images of the indicated channels of the transfected clones, stimulated or not with IS (Raji cells are labelled with a white discontinuous line), are shown. In the lower right plot, fold induction of Phospho-(Ser) PKC substrate MFI (mean+SD) in non-transfected (YFP^-^) or HA-YFP-FMNL1β-expressing (YFP-FMNL1β**^+^**), either non-stimulated (Cont) or forming synapses (IS), C3 cells, summarizing the results obtained in several (n=5) experiments. NS, not significant; **, p≤0.05. This figure is related to Fig. 6.

**Suppl. Fig. S11.**
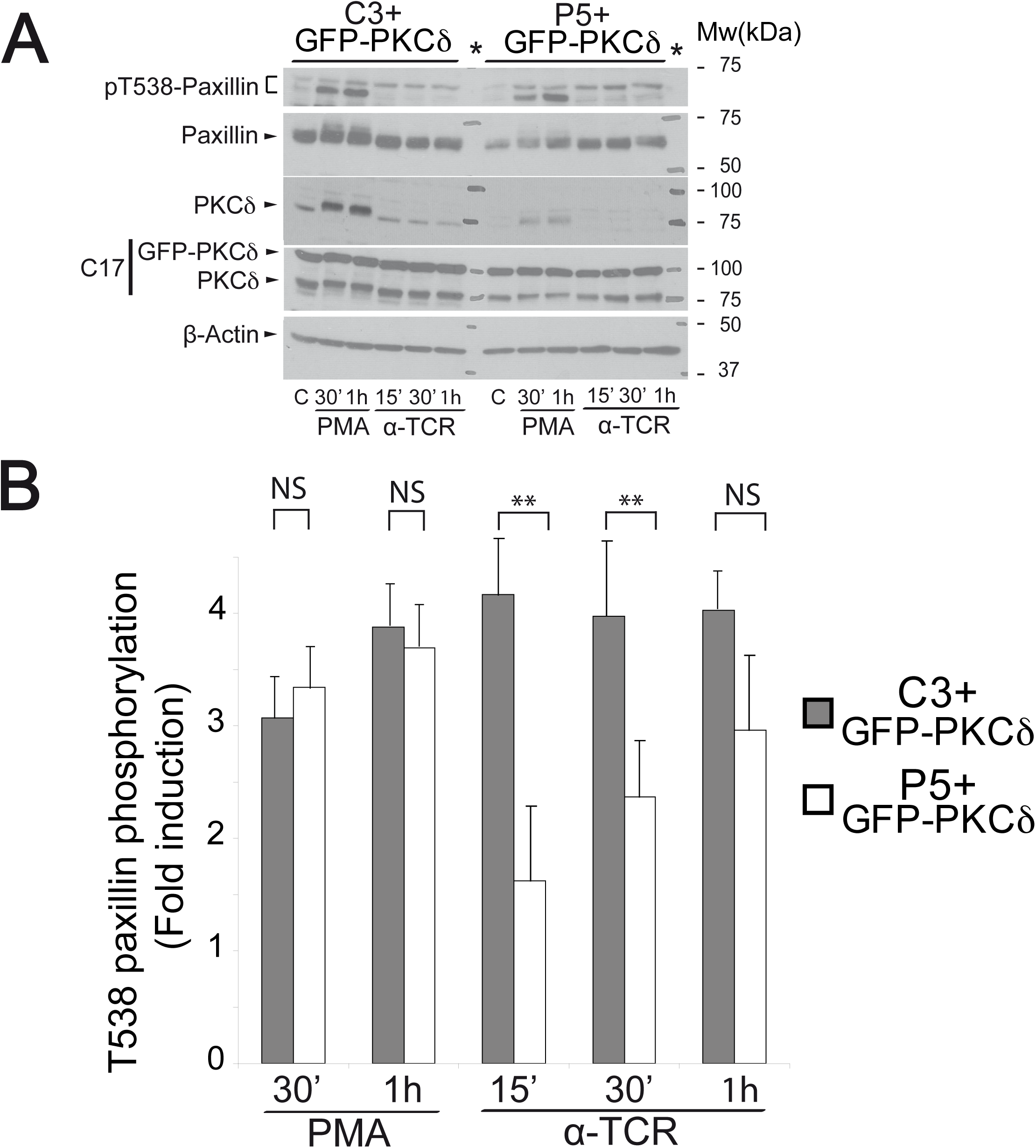
GFP-PKCδ expression partially restores T538-paxillin phosphorylation. A) C3 control and P5 PKCδ-interfered clones expressing mouse GFP-PKCδ were left untreated (C) or stimulated at the indicated times with either PMA or plastic-bound anti-TCR to induce paxillin phosphorylation. WB of cell lysates was developed with anti-phospho-T538-paxillin, anti-paxillin, anti-PKCδ, anti-ratPKCδ (C17) and anti-β-actin. The lane corresponding to the molecular weight markers is labeled (*). B) C3 control and P5 PKCδ-interfered clones expressing GFP-PKCδ were stimulated as in panel A. Subsequently, phosphorylation of paxillin at T538 was quantified by analyzing several WB similar to that described in panel A. Internal normalization of phospho-T538-paxillin signal was performed by using total paxillin signal. Data are represented as average fold induction (mean±SD) of paxillin phosphorylation in T538 relative to unstimulated C3 control or P5 PKCδ-interfered clones (*n* = 3). NS, not significant; **, p≤0.05. This figure is related to Fig. 7.

**Supplementary Video 1.** DsRed-Cent2 and CFP-CD63-expressing, C3 control clone cells were conjugated with CMAC-labelled (blue), SEE-pulsed Raji cells as described in Materials and Methods. Subsequently, the conjugates were imaged by time-lapse microscopy. In the left panel, CMAC (blue), dsRed-Cent2 (red) and CFP-CD63 (cyan) channels were merged. The middle panel corresponds to a 3x zoom of the conjugate, to properly visualize subcellular structures. In the right panel, the trajectory of the MTOC obtained by MTrackJ plugin from ImageJ (yellow line) was superimposed to the three channels (6 fps). Both the polarization of MTOC towards the IS and the simultaneous convergence of MVB towards the MTOC are evident. Results are representative of the results obtained from at least 16 time-lapse videos.

**Supplementary Video 2.** CMAC-labelled Raji cells (blue) were attached to fibronectin-coated IBIDI chamber slides and pulsed with SEE (30 min). Synaptic formation by C3 control (upper panel) and P5 PKCδ-interfered (lower panel), CFP-CD63-expressing clones, was imaged by time-lapse microscopy. In the left panels, CMAC and CFP-CD63 (cyan) channels were merged and the synaptic contacts are labelled with white arrows. In the right panels, the white lines encircle the Raji cells (blue) and the white spots label the positions of the CFP-CD63 vesicles. The trajectory of the vesicles was analysed and recorded by using the Particle Tracker plugin form Image J. The video (3 fps) shows the polarization of MVB towards the synapse contact areas occurring in the C3 control clone, but not in the P5 PKCδ-interfered clone, although both clones formed synaptic conjugates and the convergence of the vesicle trajectories was evident in both clones. One representative example is shown out of 13 synapses recorded of each clone. This video is related to Fig. 1A and suppl. Video 1.

**Supplementary Video 3.** C3 control and P5 PKCδ-interfered clones were conjugated with CMAC-labelled (blue), SEE-pulsed Raji cells. After 1 h of conjugate formation, fixed cells were labelled with phalloidin (green), and anti-CD63 (magenta) and analyzed by fluorescence microscopy using the 3D Viewer plugin of the ImageJ software, to visualize Z planes and to project the IS interface. Upper panels: merged CMAC, phalloidin and CD63 channels. Medium panels: enlarged IS area (1.8x and 2.3x zoom for C3 and P5, respectively, white rectangles in Fig. 1C) of phalloidin channel. Lower panels: enlarged IS area (1.8x and 2.3x zoom for C3 and P5, respectively, white rectangles in Fig. 1C) of merged phalloidin and CD63 channels. Frame no. 1 of the video (2 fps) corresponds to the top view shown in Fig. 1C and, after a 3D rotation, frame no. 43 corresponds to the IS interface view shown in Fig. 1C. The depletion of F-actin and the accumulation of CD63^+^ vesicles (MVB) at the synaptic central region are observed in the C3 control clone, but not in the P5 PKCδ- interfered clone. This video is related to Fig. 1C. Results are representative of the results obtained from at least 22 videos of each clone.

**Supplementary Video 4.** CMAC-labelled Raji cells (blue) were attached to fibronectin-coated IBIDI chamber slides and pulsed with SEE (30 min). Synapse formation by the C3 control clone (upper row) and the P5 PKCδ-interfered clone (lower row), expressing dsRed-Cent2 (red, yellow arrow) and previously labelled with 100 nM SirActin (green) and verapamil was imaged by wide-field, time-lapse microscopy (2 fps). In the left column, transmittance (TRANS) plus CMAC (blue) channels were merged to visualize the synaptic contact. In the second column, dsRed-Cent2 (red) plus CMAC (blue) channels were merged, whereas in the third column the three channels (CMAC, dsRed-Cent2 and SirActin) were merged. In the fourth column, the trajectory of the MTOC (encircled in white) obtained by using the MTrackJ plugin from ImageJ (yellow line) was superimposed to the three merged channels. White arrow labels synapses, yellow arrow labels MTOC, whereas magenta arrow labels plasma membrane blebbing of the Jurkat clone. Results are representative of the results obtained from at least 9 time-lapse videos of each clone. This video is related to Fig. 2D.

**Supplementary Video 5.** CMAC-labelled, SEE-pulsed Raji cells (blue) were conjugated in suspension with the C3 control clone (upper row) or the P5 PKCδ- interfered clone (lower row), expressing dsRed-Cent2 (red), that were previously labelled with 100 nM SirActin (green) and verapamil. Synaptic conjugates were imaged by confocal, time-lapse microscopy (1.5 fps), and the merged channels are shown. In the left column, transmittance (TRANS) plus CMAC (blue) channels were merged to visualize the synaptic contact. In the second column, dsRed-Cent2 (red) plus CMAC (blue) channels were merged, whereas in the third column the three channels (CMAC, dsRed-Cent2 and SirActin) were merged. Note that, in the P5 clone, but not in the C3 clone, the centrosome red signal is hardly visible, partly masked by the high centrosomal F-actin green signal, as opposed to C3. This video is related to Fig. 2D, lower panels. Results are representative of the results obtained from at least 14 time-lapse videos of each clone.

**Supplementary Video 6.** CMAC-labelled Raji cells (blue) were attached to fibronectin-coated IBIDI chamber slides and pulsed with SEE (30 min). Synapse formation by the C3 control clone (upper panels) or the P5 PKCδ-interfered clone (lower panels), co-expressing GFP-actin (green), CFP-CD63 (cyan) and pmCherry-paxillin (red), was imaged by time-lapse microscopy (2 fps). In the upper left panel for each clone, transmittance and CMAC channels were merged, whereas in the upper right panel, CMAC and GFP-actin channels were merged to visualize both the synaptic contact and the cortical F-actin burst at the IS. In the lower panels, CMAC and CFP-CD63 channels (left), or CMAC and pmCherry-paxillin channel (right) were merged to visualize the convergence/polarization of MVB and paxillin polarization to the IS, respectively. Results are representative of the results obtained from at least 17 time-lapse videos.

